# Cell type specific gene expression profiling reveals a role for the complement component C3A in neutrophil migration to tissue damage

**DOI:** 10.1101/2019.12.12.874511

**Authors:** Ruth A. Houseright, Emily E. Rosowski, Pui Ying Lam, Sebastien JM Tauzin, Oscar Mulvaney, Colin N. Dewey, David Bennin, Anna Huttenlocher

## Abstract

Following acute injury, leukocytes rapidly infiltrate into tissues. For efficient recruitment, leukocytes must sense and respond to signals from both from the damaged tissue and from one another. However, the cell type specific transcriptional changes that influence leukocyte recruitment and wound healing have not been well characterized. In this study, we performed a large-scale translating ribosome affinity purification (TRAP) and RNA sequencing screen in larval zebrafish to identify genes differentially expressed by neutrophils, macrophages, and epithelial cells in the context of wounding. We identified the complement pathway and *c3a.1*, homologous to the C3A component of human complement, as significantly increased in neutrophils in response to a wound. We report that *c3a.1*^*−/−*^ zebrafish larvae have impaired neutrophil responses to both tail wounds and localized bacterial infections, as well as increased susceptibility to infection due to a neutrophil-intrinsic function of C3A. We further show that C3A enhances migration of human primary neutrophils to IL-8 and that *c3a.1*^*−/−*^ larvae have impaired neutrophil migration *in vivo*, and a decrease in neutrophil directed migration speed early after wounding. Together, our findings suggest a role for C3A in mediating efficient neutrophil migration to damaged tissues and support the power of TRAP to identify cell-specific changes in gene expression associated with wound-associated inflammation.

## Introduction

Acute tissue injury is characterized by a rapid influx of leukocytes, both neutrophils and macrophages, into the wound microenvironment, followed by inflammatory resolution and wound healing(1). This initial recruitment of leukocytes to the wound is of critical importance; neutrophils, the most abundant cell type and the first responders to tissue damage, limit infection at the wound site (2, 3), while macrophages remove debris that would otherwise impede the repair process (4, 5).

In order for efficient recruitment to the wound to occur, leukocytes must sense and respond to a complex milieu of signals, both from the damaged tissue itself and from one another. For example, early after wounding, the damaged tissue produces a burst of hydrogen peroxide (6). This cue, originating from epithelial cells, induces signaling changes within neutrophils, including activation of the Src family kinase Lyn, which is necessary for efficient early neutrophil recruitment to the wound (7). Integration of these early wound signals is important; for example, interrupting ROS signaling for as little as 1 hour post-wounding (1hpw) can impair healing and regeneration in a larval zebrafish model 3 days later, suggesting there are transcriptional changes induced by this early signal (8).

Zebrafish represent a strong *in vivo* system to answer these questions, as they have functioning cellular and noncellular arms of the innate immune system that are largely conserved to those of humans, including neutrophils (9–11), macrophages (11–13), and the complement system (14, 15). Zebrafish have high fecundity, which increases the statistical power of experiments, and are thus an attractive model for use in large-scale genetic and drug screens. Further, zebrafish are highly amenable to live imaging of leukocyte migration in response to inflammatory stimuli (16).

We have previously reported the use of translating ribosome affinity purification (TRAP) to detect changes in gene expression in specific cell types resulting from heat shock in zebrafish (17). However, this method has not, to our knowledge, been used to detect cell type-specific differential gene expression in response to wounding. Using this method, we detected upregulation of complement system components in cells upon wounding, including c3a.

As a non-cellular arm of the innate immune system, the complement system has long been recognized to play an important role in mediating leukocyte function. As early as 1899, Paul Ehrlich recognized that immune cells express receptors that can bind the heat-labile, antimicrobial component of fresh serum, which now known to be complement (18). Most of the proteins of the complement system are synthesized by the liver in humans and circulate in blood plasma(19). However, significant quantities of complement proteins are also produced by tissue macrophages and dendritic cells (20–24), blood monocytes and neutrophils (25–30), mast cells (31, 32), and the epithelial cells of the gastrointestinal tract (33, 34), amongst others (35). The relative contribution of complement generated by each these sources has not been fully established. Further, activation of complement via either the classical, alternative, or mannose-binding lectin pathways results in a cascade of proteolytic cleavages of complement components, converging at the hydrolysis and activation of component C3 to C3A and C3B. C3B subsequently binds to other complement pathway proteins to form C5 convertase, which cleaves component C5 to C5A and C5B (19). While both C3B and C5B play important roles in antimicrobial defense, and C5A is known to be a powerful chemoattractant for neutrophils (19), C3A remains a relatively under-studied complement protein, and its specific role in leukocyte recruitment in the absence of infection is unclear.

The complement system is evolutionarily old, with elements of the cascade present in species from protostomes to mammals (36). Zebrafish express complement proteins highly conserved to human C3A and C3B (37); however, unlike humans, zebrafish express multiple forms of C3 that are encoded on different genes (14, 15). While in humans, the full-length complement component C3 is cleaved during complement activation to produce C3A and C3B (19), zebrafish C3A and C3B are encoded on different genetic loci, and indeed, different chromosomes (38). The possibility of manipulating C3A expression at the genetic level, independent of C3B expression, without disrupting the expression of important downstream complement components such as C5A, makes the zebrafish an attractive model for studying the specific effects C3A on leukocyte responses to tissue damage.

In this work, we report that TRAP-RNAseq of larval zebrafish identifies genes differentially expressed in neutrophils, macrophages, and epithelial cells in response to wounds. Our data identify upregulation of the complement pathway in all cell types, with specific, statistically significant upregulation of *c3a.1*, homologous to the C3A component of human complement, in neutrophils. We find that *c3a.1* plays an important role in neutrophil recruitment, as mutation of *c3a.1* results in impaired neutrophil recruitment to wounds. Neutrophil recruitment to and survival of localized bacterial infections is also impaired in *c3a.1*^*−/−*^ larvae. We find that these defects in neutrophil recruitment are likely due to decreased neutrophil migratory speed in the early post-wounding period. We further show that, *in vitro*, C3A does not serve as a direct chemoattractant for human primary neutrophils, but instead sensitizes neutrophils to respond to IL-8, suggesting a role for C3A in priming neutrophils to respond to other inflammatory cues.

## Methods

### Zebrafish lines, maintenance, and genotyping

All zebrafish were maintained according to protocols approved by the University of Wisconsin-Madison Institutional Animal Care and Use Committee, as described previously(39). Previously published zebrafish lines were used (**Supplemental Table 1**). Larvae were anesthetized using 0.2 mg/mL tricaine before any experimentation. Zebrafish containing the mutant *c3a.1* allele sa31241 were isolated through the Sanger Zebrafish Mutation Project, Wellcome Sanger Institute, and obtained from the Zebrafish International Resource Center (ZIRC). This allele will be referred to as *c3a.1*^*−/−*^ herein. The sa31241 point mutation was detected by restriction fragment length polymorphism (RFLP) analysis. DNA was isolated in 50mM NaOH, the mutated region amplified with GoTaq (Promega) (**Supplemental Table 2**), and a restriction enzyme (DraI, NEB) directly targeting the mutant copy, but not the wild-type copy, was directly added with buffer. Digests were incubated overnight and run on a 2% agarose gel to evaluate the presence of mutant and/or wild-type bands.

### Purification of mRNA from TRAP zebrafish larvae and RNA sequencing

3 dpf *Tg (lyz:EGFP-L10a), Tg(mpeg1:EGFP-L10a)*, or *Tg(krt4:EGFP-L10a)* (17) zebrafish larvae were anesthetized using 0.2 mg/mL tricaine and subjected to multiple tail fin wounds using a 33 gauge needle (Fig. 1A). A previously-published protocol for TRAP mRNA purification from zebrafish (17) was performed, with slight modifications. Briefly, a QIAshredder (Quiagen) was used to homogenize the larvae prior to immunoprecipitation. mRNA was isolated from immunoprecipitated polysomes from 50 pooled wounded fish or unwounded controls using TRIzol reagent (Invitrogen). 4 paired replicates were collected. RNA quality and concentration was assessed on Agilent RNA PicoChip and samples with a concentration <200 pg/µl were concentrated in a SpeedVac. As RNA concentrations were low, library prep was done with the NuGEN Ovation Single Cell RNA-Seq System with 20 cycles of amplification. Libraries were then checked by QuBit and an AATI Fragment Analyzer for concentration and fragment size. Adaptors with barcodes were used and samples were sequenced on an Illumina HiSeq with an average of 8 samples per lane. Sequencing generated 27.5 million single end reads per sample on average. Gene-level read counts were estimated using RSEM v1.2.20(40) and Bowtie v1.1.1 (41) with the Ensembl v83 annotation of the GRCz10 assembly of the zebrafish genome. One *Tg (lyz:EGFP-L10a)* wounded sample was removed from the analysis during quality control assessments because it clustered most closely with other samples sequenced at the same time that had been generated via a different protocol.

**FIGURE 1.**
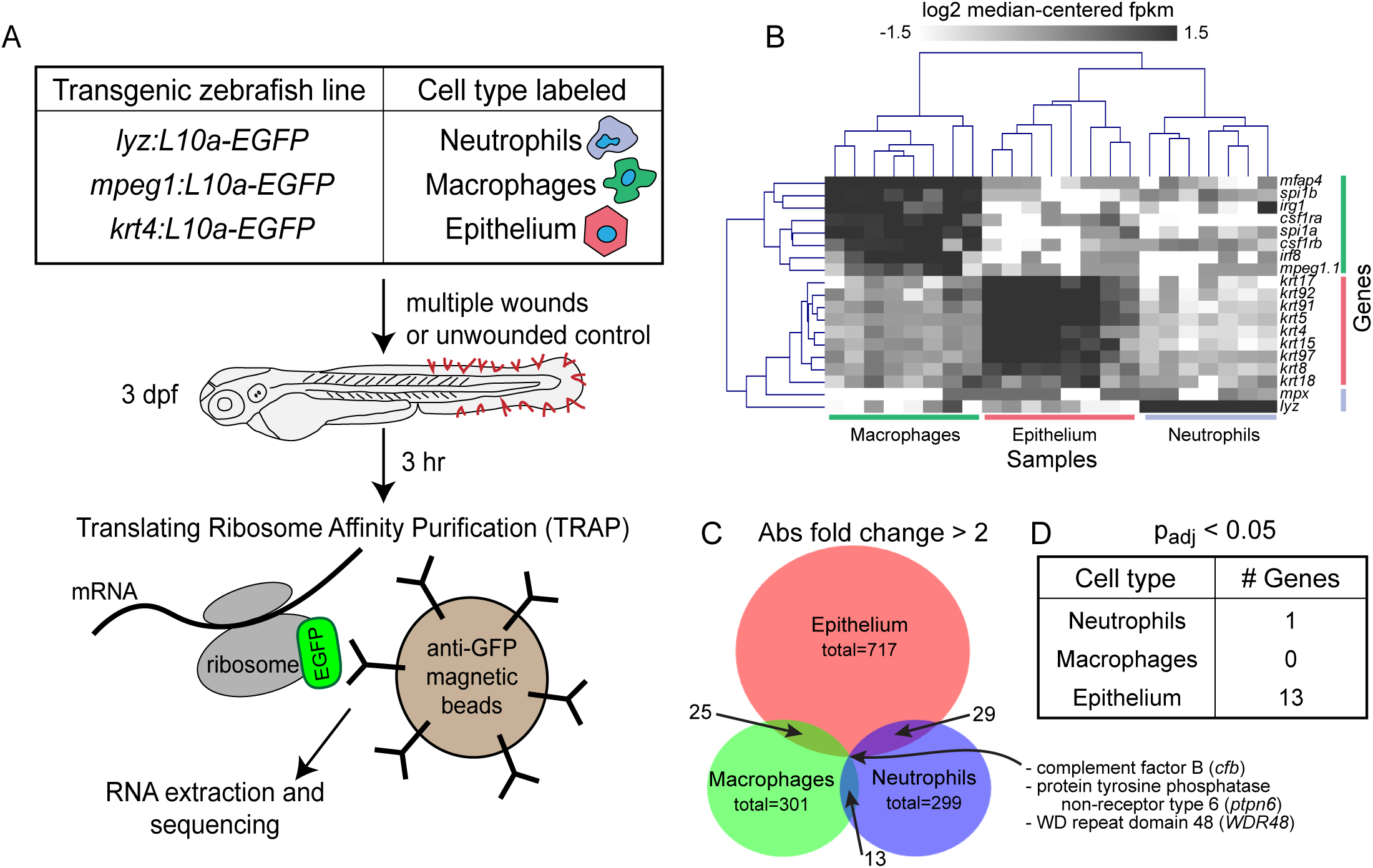
TRAP-RNAseq identifies differential expression of genes by neutrophils, macrophages, and epithelial cells in response to wounding. (A) Experimental setup. 3 dpf transgenic zebrafish larvae expressing an EGFP-tagged copy of the ribosomal subunit L10a specifically in neutrophils (*lyz*), macrophages (*mpeg1*), or epithelium (*krt4*) were subjected to multiple fin tissue wounds. 3 hours later, larval tissue was homogenized and ribosomes were isolated with α-EFP immunoprecipitation. RNA was then extracted and subject to Illumina sequencing. (B) Expression of a priori tissue-specific genes across all analyzed samples. Columns represent samples, labeled by cell type-specific promoter used, rows represent known cell-type-specific genes. (C) Venn diagram of genes found to be more than 2-fold changed by wounding in each cell type. (D) Genes with expression significantly altered (p_adj_ < 0.05) upon wounding.

Differentially expressed genes identified by RNA-seq were called using the DESeq2 R package (42). The design formula for the generalized linear model used with DESeq2 was “~ replicate + condition” where “condition” was the combination of cell type and treatment for each sample. Statistical testing for differential expression within each cell type was performed using the Wald test implemented in the DESeq2 package. Translating RNAs with at least a 2-fold change in their relative abundance with a Benjamini-Hochberg corrected *P* value (FDR) ≤ 0.05 were considered statistically significant.

Human homologs of zebrafish genes were extracted from Ensembl using the BioMart tool. Gene Set Enrichment Analysis(43, 44) was performed by comparing gene expression data mapped to these human homologs to Hallmark gene sets (v6.2) from the Molecular Signatures Database (Broad Institute)(45). The gsea3 java release was run using all default settings. Heatmaps were generated with Multiple Experiment Viewer (MeV) and Venn diagrams were generated and overlaps determined by BioVenn(46).

### RT-qPCR

RNA was extracted from approximately 50 pooled, 3 dpf *c3a.1*^*+/+*^ or *c3a.1*^*−/−*^ larvae using TRIzol reagent (Invitrogen). cDNA was then synthesized using SuperScript III RT and oligo-dT (Invitrogen). Using this cDNA as a template, quantitative PCR (qPCR) with FastStart Essential DNA Green Master (Roche) and a LightCycler96 (Roche) was performed. Fold changes in gene expression over control conditions, normalized to *ef1a*, were calculated from Cq values (47). Primers used to amplify *c3a* orthologues(48), and *ef1a*(49) have been described previously. Due to high identity percentage, *c3a.2-3* and *c3a.7-8* were amplified and analyzed together, as previously described (48). Primer sequences used in this study can be found in **Supplemental Table 2**.

### Tail transection

*C3a.1*^*+/−*^ adults were in-crossed. 3 dpf larvae were wounded by tail transection using a no. 10 Feather surgical blade. To visualize neutrophils in the wound microenvironment, the larvae were fixed at 2 hpw or 8 hpw in 4% paraformaldehyde in 1X PBS overnight at 4°C. Sudan Black B staining was performed as described previously (50). Fixed larvae were imaged using a zoomscope (EMS3/SyCoP3; Zeiss; Plan-NeoFluar Z objective) and then genotyped as above. For macrophage quantification, *c3a.1*^*+/−*^ adults carrying a *mpeg1:GFP* transgene (13) were in-crossed. At 3 dpf, larvae were pre-screened for fluorescence on a zoomscope. Tail wounding was then performed as described above and the larvae were fixed in 1.5% formaldehyde overnight at 4°C. Fixed larvae were imaged using a zoomscope and genotyped as above. All image analysis was performed using Zen 2012 (blue edition, Carl Zeiss), blinded to genotype.

### *Pseudomonas* infections

3 dpf *c3a.1*^*+/+*^ or ^*−/−*^ larvae on a WT AB or *Tg(mpx:mcherry-2A-rac2*^*D57N*^) background (51) were infected with *P. aeruginosa* PAK (pMF230, expresses GFP) as previously described (52, 53). PAK (pMF230) was a gift of Dara Frank (Medical College of Wisconsin). A single colony was inoculated overnight in LB. In the morning, the culture was diluted 1:5 and grown for an additional 1.25 hours. The OD was measured (600nm). The final inoculum was prepared by pelleting the bacterial suspension by centrifugation and resuspending the bacteria to achieve the desired bacterial density in 1X PBS containing 10% glycerol and 2% PVP-40 (to prevent needle clogging). Phenol red dye was added at a final concentration of 0.5% to visualize success of the injection. To monitor CFUs, the injection product was plated on LB and incubated overnight. Injected CFUs are noted in the figure legends. For survival experiments, infected larvae were placed into individual wells of a 96 well plate and survival was monitored daily for 5 dpi. For neutrophil recruitment experiments, larvae were fixed at 1 hpi or 6 hpi in 4% paraformaldehyde in 1X PBS overnight at 4°C. Sudan Black B staining was performed as described previously (50), and injection success was further confirmed by visualization of GFP-positive bacteria in the otic vesicle on a spinning disk confocal microscope (CSU-X, Yokogawa) as described below, without mounting in agarose. Imaging of the otic vesicle region for neutrophil enumeration was performed using a zoomscope (EMS3/SyCoP3; Zeiss; Plan-NeoFluar Z objective). Image analysis was performed using Zen 2012 (blue edition, Carl Zeiss).

### Photoconversion

Adult *c3a.1*^*+/−*^ zebrafish carrying an *mpx:Dendra2* transgene (54) were in-crossed and embryos collected and incubated to 3 dpf. Larvae were prescreened for fluorescence using a zoomscope (EMS3/SyCoP3; Zeiss; Plan-NeoFluar Z objective) and mounted in ZWEDGI devices as previously described (55). An imaging sequence was performed for each larva comprising an initial series of 2 overlapping Z-stacks of the region of the caudal hematopoietic tissue (CHT) and photoconversion of the neutrophils within the CHT. This was followed by a second series of 2 overlapping Z-stacks to confirm that photoconversion occurred. Photoconversion was performed using a laser scanning confocal microscope (FluoView FV1000; Olympus) with numerical aperature (NA) 0.75/20X objective. The following stimulation settings were used: 40% 405 nm laser transmissivity, 10 µs/pixel dwell time, and 45 second total stimulation time. Larvae were removed from the ZWEDGI devices following photoconversion and subjected to wounding by tail transection as above. Larvae were subsequently imaged live at 3 hpw using a spinning disk confocal microscope as described below and then genotyped as above. Image analysis was performed using Zen 2012 (blue edition, Carl Zeiss), blinded to genotype.

### Live imaging and image quantification

3 dpf *c3a.1*^*+/+*^ or ^*−/−*^ larvae carrying a *mpx:mcherry* transgene (56) were pre-screened for fluorescence on a zoomscope (EMS3/SyCoP3; Zeiss; Plan-NeoFluar Z objective). For imaging over 1-3 hours, larvae were mounted in ZWEDGI devices as previously described (55) and retained in place using 2% low melting point agarose applied to the head. Images were acquired every 3 minutes using a spinning disk confocal microscope (CSU-X, Yokogawa, NA 0.3/10X EC Plan-NeoFluar objective) with a confocal scanhead on a Zeiss Observer Z.1 inverted microscope equipped with a Photometrics Evolve EMCCD camera. Each image comprised a 50 µm z-stack, with 11 slices taken at 5 µm intervals. Images were analyzed and maximum intensity projections were made using Zen 2012 (blue edition) software (Carl Zeiss). To track cell motility, time series were analyzed in Imaris (Bitplane) and neutrophil mean track speed, track displacement, and track straightness, as well as instantaneous velocity for each neutrophil at each point in the time series, were calculated using the “spots” tool as previously described (57). To count total neutrophils and quantify neutrophil distribution in photoconversion experiments, 12 overlapping images were acquired to capture the full length and width of each larva and image analysis and neutrophil counts were performed using the “events” tool in Zen 2012 (blue edition, Carl Zeiss).

### Human primary neutrophil purification

Peripheral blood neutrophils from human blood were purified using the Miltenyi Biotech MACSxpress Neutrophil Isolation Kit according to the manufacturer’s instructions (Miltenyi Biotec, #130-104-434) and residual red blood cells were lysed using MACSxpress Erythrocyte Depletion Kit (Miltenyi Biotec, #130-098-196) All donors were healthy and informed consent was obtained at the time of the blood draw according to the requirements of the institutional review board (IRB).

### Chemotaxis Assay

Chemotaxis was assessed using a microfluidic device as described previously (58). In brief, polydimethylsiloxane devices were plasma treated and adhered to glass coverslips. Devices were coated with 10 μg/mL fibrinogen (Sigma) in PBS for 30 min at 37°C, 5% CO_2_. The devices were blocked with 2% BSA-PBS for 30 min at 37°C, 5% CO_2_, to block non-specific binding, and then washed twice with mHBSS. Cells were stained with calcein AM (Molecular Probes) in PBS for 10 min at room temperature followed by resuspension in modified Hank’s balanced salt solution (mHBSS). Cells were seeded at 5 × 10^6^/mL to allow adherence for 30 min before addition of chemoattractant. Either 3μM Complement Component C3a (R&D Systems #3677-C3-025) or 3μM Complement Component C5a (R&D Systems #2037-C5-025/CF) or 1 μM IL-8 (R&D Systems # 208-IL-10/CF) was loaded onto the devices. For pretreated samples, cells were incubated in 3μM C3a or blank mHBSS for 30min before seeding device. Cells were imaged for 45 min every 30 s on a Nikon Eclipse TE300 inverted fluorescent microscope with a 10× objective and an automated stage using MetaMorph software (Molecular Devices). Automated cell tracking analysis was done using JEX software (59) to calculate chemotactic index and velocity.

### Statistical analyses

For neutrophil quantification and migration analyses, 3-4 independent replicate experiments were performed. Replicate numbers are noted in the figure legends. Experimental conditions were compared using analysis of variance. The results were summarized and plotted in terms of least squares adjusted means and standard errors.

For survival curves, 3 independent experiments were performed. Results were pooled and analyzed by Cox proportional hazard regression analysis, with experimental conditions included as group variables. Statistical analyses were performed using R version 3.4.4 and graphical representations were made using GraphPad Prism version 7. Significance was defined as *P* < 0.05. The resulting *P* values are included in the figure legends for each experiment.

For quantification of neutrophil instantaneous speed over time, a linear mixed effect regression model was used. Genotype and time were treated as fixed effects, with experimental replicate, fish, and neutrophil (within fish) treated as random effects. Statistical analyses were performed in R version 3.5.1, using the associated lme4 package. Reported *P* values are 2-sided and level of statistical significance preset to 0.05, with no adjustment for multiplicity.

## Results

### TRAP-RNAseq identifies genes differentially regulated in neutrophils, macrophages, and epithelial cells in response to wounding

Communication between multiple cell types, including leukocytes and epithelial cells, is essential to allow cells of the innate immune system to effectively navigate complex interstitial tissues to reach the wound microenvironment (60). However, the relative transcriptional contributions of each cell type are incompletely understood. To identify cell-specific signals that are differentially expressed in different cell types in response to wounding, we performed a large-scale translating protein affinity purification and RNA sequencing (TRAP-RNAseq) screen (Fig. 1A). Briefly, 3 dpf transgenic zebrafish larvae expressing an EGFP-tagged copy of the ribosomal subunit L10a specifically in neutrophils, macrophages, or epithelium were subjected to multiple fin tissue wounds. 3 hours later, larval tissue was homogenized and ribosomes were isolated with α-GFP immunoprecipitation. RNA was then extracted. Illumina sequencing confirmed expression of *a priori*-selected, known cell-type-specific genes across all analyzed samples, validating our method (Fig. 1B). We then focused our analysis solely on zebrafish genes that have identified human homologs. From these genes, 299 were identified to be at least 2-fold changed (upregulated or downregulated) in neutrophils, 301 in macrophages, and 717 in epithelial cells. In neutrophils, we were surprised to find that only a single gene was statistically significantly upregulated in response to wounding: *c3a.1* (Fig. 1D). Although other changes in gene expression did not reach statistical significance, which is likely due to high variability among samples and small numbers of replicates performed, we expect that 2-fold differential expression is potentially biologically relevant. It is interesting to note that relatively few genes were more than 2-fold differentially expressed in more than one cell type (Fig. 1C **and Supplemental Table 3**).

### TRAP-RNAseq identifies the complement pathway and *c3a.1* as factors upregulated upon wounding

C3a.1 shares an approximately 43% amino acid similarity to the human C3a component of complement (48). Gene set enrichment analysis (GSEA) of Hallmark genes from the Molecular Signatures Database to identify groups of genes sharing a common biologic function (43) further showed enrichment of genes involved in the complement pathway in wounded fish, compared with unwounded controls (Fig. 2A). Although only *c3a.1* showed a statistically significant increase in mRNA expression in neutrophils following wounding, other complement pathway components, including *c5* and *c9*, also showed trends toward increased expression in neutrophils (Fig. 2B). Non-significant increases in *c3a.1, c5* and *c9* expression were also evident in macrophages. Further, complement factor B (*cfb*) was one of only 3 genes that were differentially expressed in all 3 cell types (Fig. 1C). Taken together, these data suggest an important role for the complement pathway in general, and *c3a.1* in particular, in orchestrating innate immune responses in the context of wounding.

**FIGURE 2.**
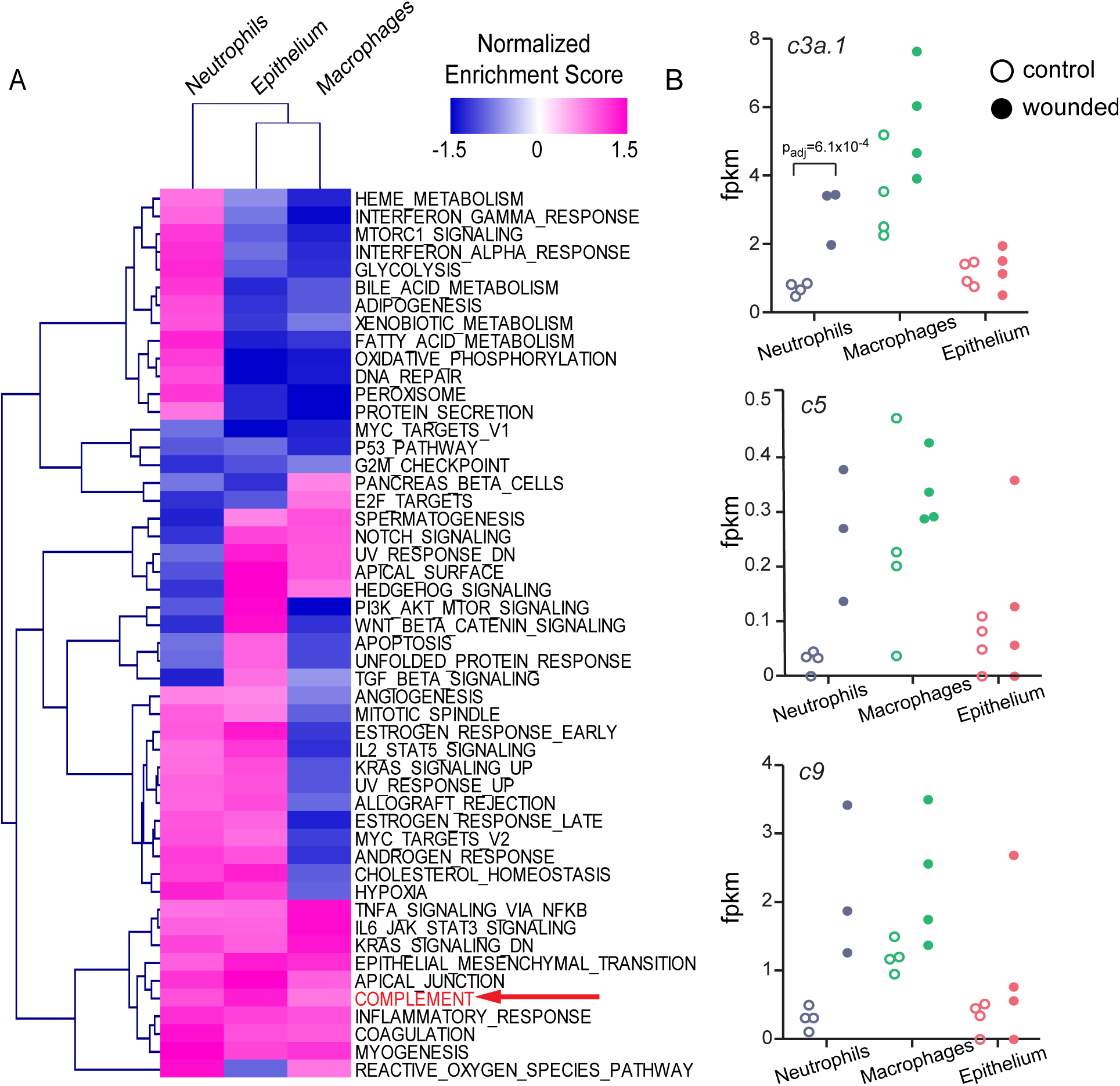
TRAP-RNAseq identifies upregulation of the complement pathway and c3a.1 in response to wounding. (A) Normalized enrichment scores of Molecular Signatures Database Hallmark Gene Sets (rows) in each cell type (columns), from Gene Set Enrichment Analysis (GSEA). (B) Expression (fpkm) of three complement-pathway genes (*c3a.1*, *c5*, and *c9*) across all three cell types. Each dot represents one replicate.

### Validation of *c3a.1*-deficient zebrafish lines

In order to investigate the role of *c3a.1* in leukocyte responses in the context of wounding, we obtained zebrafish expressing an A to T nonsense mutation in exon 22 of 41 of the *c3a.1* sequence, producing a premature stop codon (sa31241, Sanger) (61) (Fig. 3A). This premature stop codon occurs prior to the predicted thioester bond and α2 macroglobulin-complement domains of the C3A protein that characterize an anaphylatoxin (62). qPCR of cDNA from pooled 3 dpf *c3a.1*^*−/−*^ larvae confirmed loss of *c3a.1* mRNA, compared with *c3a.1*^*+/+*^ controls, and showed no significant compensatory upregulation of the other *c3a* orthologues expressed at this stage of larval development (Fig. 3B). Amplification of *c3a.2-3, c3a.4*, and *c3a.5* by RT-qPCR was not observed; this is in agreement with existing reports that these orthologues are not expressed in 3 dpf larvae (48). Expression of other major complement pathway genes based on qPCR of cDNA from pooled 3 dpf *c3a.1*^*−/−*^ larvae was similar to that of *c3a.1*^*+/+*^ controls: although a modest, non-significant decrease in *c3b.1* mRNA was noted, *c3b.2* mRNA was unchanged and a significant, potentially compensatory, increase in *c5a* mRNA was observed (Fig. 3B). *c3a.1*^*−/−*^ larvae hatched in normal Mendelian ratios from *c3a.1*^*+/−*^ in-crosses; however, the tail length of *c3a.1*^*−/−*^ larvae was significantly shorter than *c3a.1*^*+/+*^ clutch-mates at 4 dpf (**Fig. S1**). Heterozygotes were indistinguishable from their wild-type clutch-mates (data not shown).

**FIGURE 3.**
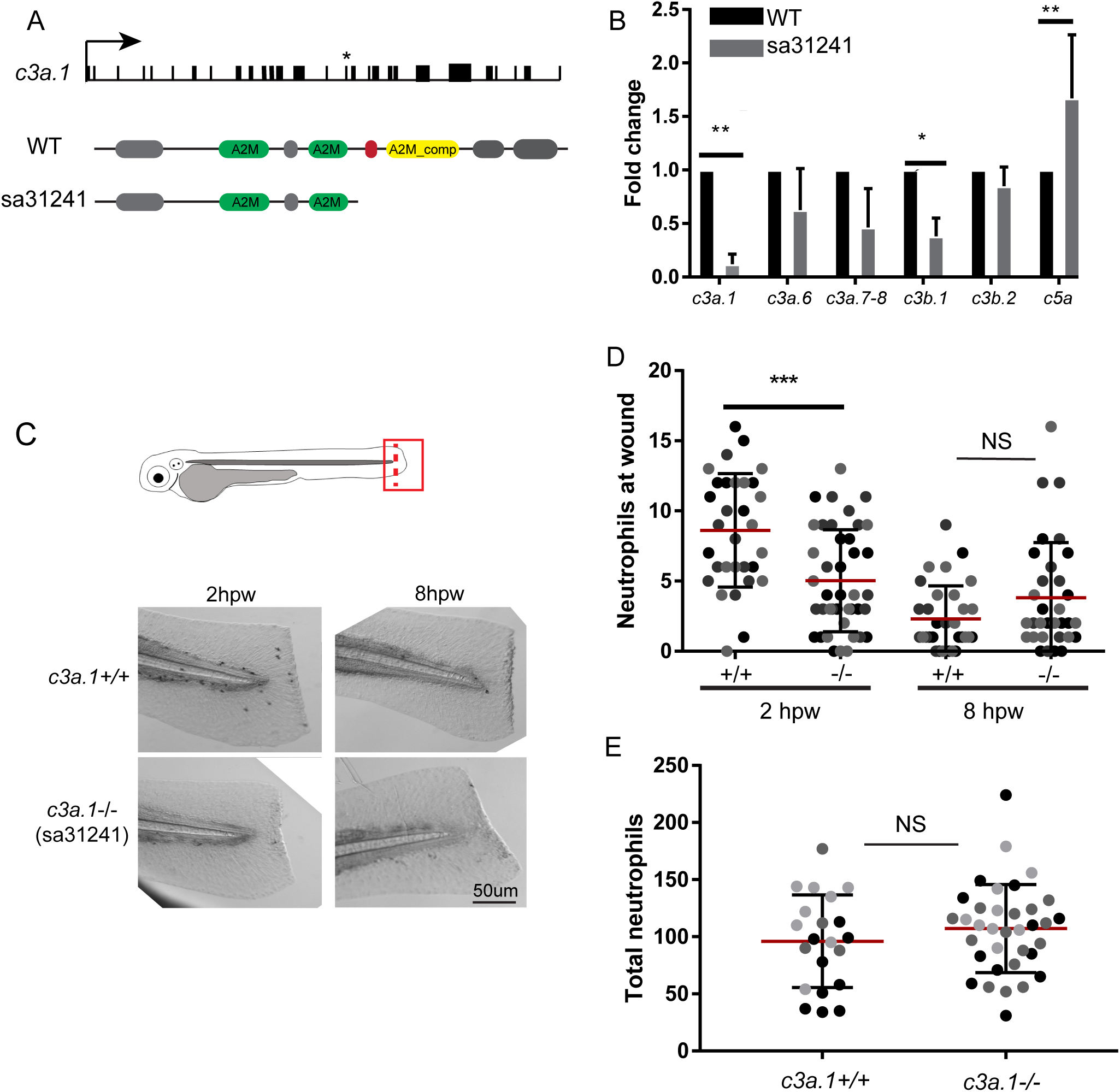
Global depletion of *c3a.1* decreases early neutrophil recruitment to a wound. (A) Schematic of *c3a.1* locus (top), with exons represented as black boxes. * indicates approximate location of A>T nonsense mutation in exon 22 in sa31241 mutant. Schematic of WT (middle) and mutated (bottom) C3a.1 protein, with selected *Pfam-*predicted domains noted. Green: α2 macroglobulin; red: thiolester bond-forming region; yellow: α2 macroglobulin complement component. (B) RT-qPCR validation of *c3a*, *c3b*, and *c5a* orthologue expression in pooled WT and *c3a.1*^*−/−*^ (sa31241) whole zebrafish larvae, normalized to WT expression for each gene and to *ef1α*, with data expressed as mean +/− SEM. Data comprise 3 independent experiments, performed in triplicate, n = 50 larvae per condition per experiment. B, C. 3 dpf WT or *c3a.1*^*−/−*^ zebrafish larvae were subjected to wounding by tail transection (dashed line), subsequently stained with Sudan Black B, and the tail region (box) was imaged. (B) Representative images and (C) quantification of neutrophil recruitment following tail transection in WT (n = 31, 2 hpw; 32, 8 hpw) and *c3a.1−/−* (n = 41, 2 hpw; 33, 8 hpw) larvae, with data expressed as mean +/− SEM. (D) Quantification of total neutrophil counts in WT (n = 21) and *c3a.1*^*−/−*^ (n = 35) larvae, with data expressed as mean +/− SEM. For C and D, each dot represents one larva; colors represent results of 3 independent experiments. *p<0.05, **p<0.01, ***p<0.001

### Global depletion of *c3a.1* decreases neutrophil recruitment to wounds

Because *c3a.1* expression is significantly increased in neutrophils in response to wounding, we first investigated the neutrophil response to wounding in *c3a.1*^*−/−*^ larvae. Compared with *c3a.1*^*+/+*^ controls, *c3a.1*^*−/−*^ larvae had significantly decreased numbers of neutrophils in the wound microenvironment at 2 hpw (Fig. 3C-D). However, by 8 hpw, neutrophil numbers at the wound did not differ between *c3a.1*^*−/−*^ and *c3a.1*^*+/+*^ larvae (Fig. 3C-D), suggesting that the neutrophil recruitment phenotype induced by *c3a.1* depletion is confined to the early post-wounding period. Changes in neutrophil numbers at the wound are due to a specific change in the recruitment response, as whole-larvae total neutrophil numbers did not differ between *c3a.1*^*−/−*^ larvae and *c3a.1*^*+/+*^ fish (Fig. 3E). These findings are in line with a neutrophil recruitment phenotype reported by Forn-Cuni, et al., in response to *c3a.1* knockdown using antisense morpholino (48). Macrophage recruitment to tail transection wounds did not differ between *c3a.1*^*−/−*^ and *c3a.1*^*+/+*^ larvae (**Fig. S2**). In addition, *c3a.1*^*−/−*^ larvae displayed significantly decreased regenerate fin length at 24, 48, and 72 hours post-wounding, compared with *c3a.1*^*+/+*^ clutchmates (**Fig. S3**). Taken together, these findings suggest that C3A modulates neutrophil wound responses and wound healing in zebrafish larvae.

### Global depletion of *c3a.1* decreases neutrophil recruitment to, and survival of, localized bacterial infection

The complement system plays an essential role in pathogen recognition and clearance, and patients with deficiencies in the classical complement pathway are susceptible to pyogenic infections (63). Thus, we next measured the ability of neutrophils in *c3a.1-*deficient larvae to migrate to localized bacterial infections. Using an established model of localized *Pseudomonas aeruginosa* infection of the otic vesicle (64), we found that *c3a.1*^*−/−*^ larvae had fewer neutrophils at the site of the infection at both 1 hour post-infection (hpi) and 6 hpi, compared with *c3a.1*^*+/+*^ controls (Fig. 4 A-B). Neutrophils are believed to be the main cell type responsible for resistance to *Pseudomonas* infections (65). Consistent with the defect in neutrophil recruitment we observed, we found that *c3a.1*^*−/−*^ larvae had increased susceptibility to *Pseudomonas* infection, with ~50% of infected larvae dying by only 1 dpi. In contrast, >95% of *c3a.1*^*+/+*^ larvae survived to 5 dpi (hazard ratio: *c3a.1*^*−/−*^ vs. *c3a.1*^*+/+*^ = 11.091) (Fig. 4C).

**FIGURE 4.**
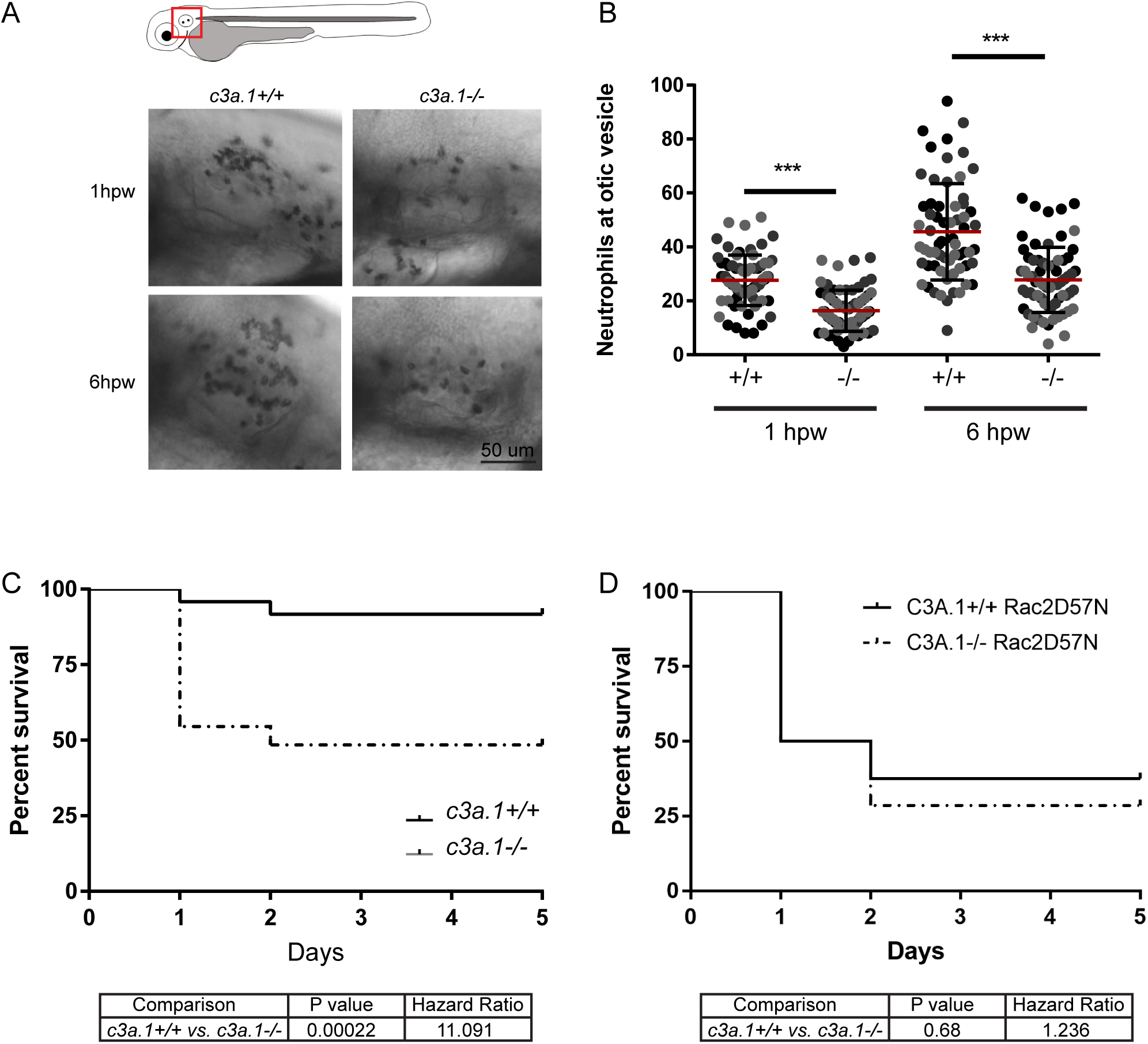
Global depletion of *c3a.1* decreases neutrophil recruitment to, and subsequent survival of, localized bacterial infection. A, B. WT (n = 71, 1 hpw; 65, 6 hpw) and *c3a.1*^*−/−*^ (n = 71, 1 hpw; 68, 6 hpw) larvae were inoculated with 1000 CFU *Pseudomonas aeruginosa* in the left otic vesicle and subsequently stained with Sudan Black B. (A) Representative images and (B) quantification of neutrophil recruitment following otic vesicle infection, with data expressed as mean +/− SEM. Each dot represents one larva; colors represent results of 3 independent experiments. ***p<0.001 (C) WT (n = 24) and *c3a.1*^*−/−*^ (n = 32) larvae were infected with 7500 CFU *Pseudomonas aeruginosa* in the left otic vesicle and survival was tracked over 5 days post-infection. (D) To test whether survival was neutrophil-dependent, *c3a.1*^*+/+*^ (n = 16) and *c3a.1−/−* (n = 14) larvae with neutrophils that are mcherry-labeled and carry a mutation in *rac2* rendering them migration-defective (*Tg*(*mpx:rac2*^*D57N*^*-mcherry*)) were infected with *Pseudomonas aeruginosa* as in C and survival was tracked over 5 days post-infection. C and D each comprise 3 independent experiments.

### *C3a.1* mediates resistance to *Pseudomonas aeruginosa* infection in a neutrophil-dependent manner

Increased susceptibility to localized *Pseudomonas* infection in *c3a.1*^*−/−*^ larvae could be due to impaired neutrophil recruitment or function, and/or the loss of other complement-mediated effects since C3A is known to have potent antimicrobial activity(66). To determine whether increased susceptibility to *Pseudomonas* infection in *c3a.1*^*−/−*^ larvae is due to neutrophil-intrinsic activity, we crossed the *c3a.1*-deficient line to the *Tg(mpx:mcherry-2A-rac2^D57N^)* line, in which mcherry-labeled neutrophils express a dominant negative form of Rac2 and are thus rendered migration-deficient. As we have previously reported (51), *c3a.1*^*+/+*^ larvae with neutrophils expressing *rac2*^*D57N*^ have increased susceptibility to *Pseudomonas* infection, with ~50% mortality at 1 dpi. In comparison, we noted no significant change in susceptibility in *c3a.1*^*−/−*^ larvae with neutrophils expressing *rac2*^*D57N*^, compared with *c3a.1-* intact *rac2*^*D57N*^ larvae (HR *c3a.1*^*−/−*^ *rac2*^*D57N*^ vs. *c3a.1*^*+/+*^ *rac2*^*D57N*^ = 1.236) (Fig. 4D). Therefore, the increase in susceptibility to *Pseudomonas* infection we observed in *c3a.1*^*−/−*^ larvae expressing wild-type *rac2* is predominantly due to a neutrophil-intrinsic function of c3a, possibly due to the reduction in numbers of neutrophils found at the infection site.

### Depletion of *c3a.1* does not alter neutrophil egress from hematopoietic tissues following wounding

We next asked how C3A controls neutrophil mobilization from hematopoietic tissue. In response to inflammatory signals, zebrafish neutrophils may be released directly from hematopoietic tissues or recruited from a population of neutrophils already patrolling in peripheral tissues (67). C3a has been shown in mice to help to retain immature neutrophils in hematopoietic tissues (68, 69). C3 and C3A receptor-deficient mice subsequently have faster and more pronounced neutrophil egress from bone marrow in response to inflammatory stimuli (70). Although this finding is opposite to the decreased neutrophil numbers that we observe at a wound in larval zebrafish (Fig. 1C-D), we still wanted to determine whether decreased neutrophil numbers at inflammatory sites in *c3a.1*^*−/−*^ larvae were due to a difference in neutrophil recruitment from hematopoietic tissue. At 3 dpf, the primary organ of hematopoiesis in the larval zebrafish is the caudal hematopoietic tissue (CHT), in which hematopoiesis resembles that within the mammalian fetal liver (71). We crossed the *c3a.1*-deficient line to the *Tg(mpx:dendra2*) line, in which neutrophils are labeled with the photoconvertible fluorophore Dendra2, enabling fate tracking of neutrophils originating from the CHT over time (54). We photoconverted neutrophils in the CHT and then subjected the larvae to tail transection. We subsequently counted both the photoconverted neutrophils remaining in the CHT and those mobilized to the periphery at 3 hpw (**Fig. S4A**). Neither the number of neutrophils retained in the CHT nor the number mobilized neutrophils differed between *c3a.1*^*+/+*^ and *c3a.1*^*−/−*^ larvae (**Fig. S4B-C**). This suggests that decreased neutrophil numbers at the wound in *c3a.1*^*−/−*^ larvae are not due to alterations in neutrophil egress from the hematopoietic tissue and led us to more closely examine neutrophil interstitial migration.

### C3A is not a direct chemoattractant for human primary neutrophils *in vitro*, but may sensitize neutrophils to respond to other chemoattractants

C5A has been well-characterized as a potent neutrophil chemoattractant (72). C3A and C5A are highly structurally similar and share a 36% amino acid identity (73); however, the effects of C3A on neutrophil polarization and migration are less well understood. Numerous authors have reported that C3A does not serve as a neutrophil chemoattractant *in vitro* (72, 74, 75). Using *in vitro* microfluidic systems, we first confirmed that primary human neutrophils show strong directional migration toward an established gradient of C5A, but do not migrate toward an established gradient of C3A (Fig. 5A-B **and Movie 1**).

**FIGURE 5.**
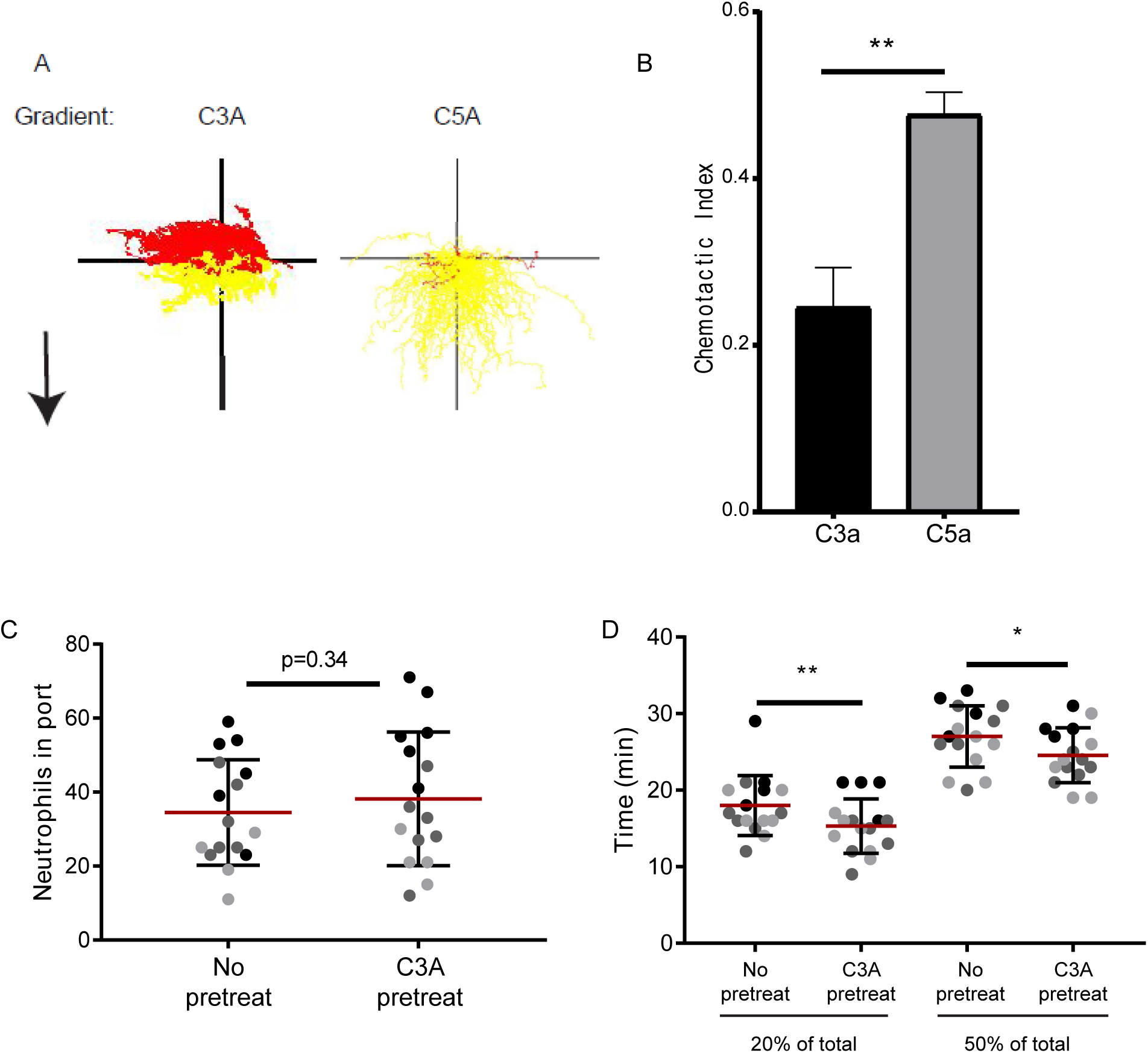
C3A sensitizes human primary neutrophils to IL-8 in vitro. (A)Representative tracks of 5 × 10^6^ human neutrophils exposed to a gradient of either C3A or C5A in a microfluidic device. Yellow tracks represent net migration toward source, and red tracks represent net migration away from the source. Black arrow indicates direction of gradient from low concentration to high concentration. (B) Quantification of chemotactic index of human neutrophils exposed to a gradient of either C3A (3 µM source) or C5A (3 µM source), expressed as mean+/−SEM. **p<0.01. (C-D) Human primary neutrophils were pre-treated with 3 µM C3A for 30 min and their migration characteristics toward a gradient of IL-8 (1 uM source), compared with non-pretreated neutrophils. (C) Quantification of total number of neutrophils reaching the chemotactic source in 45 minutes, expressed as mean +/− SEM. (D) Quantification of time required for 20% (left) or 50% (right) of all neutrophils that will reach the chemotactic source over 45 minutes to arrive, expressed as mean +/− SEM. For C and D, each dot represents the result of one technical replicate; colors indicate the results of 3 biologic replicates. *p<0.05, **p<0.01.

C3A has, however, been implicated in enhancing the homing responses of both hematopoietic progenitor cells and B cells to CXCL-12 (SDF-1) (69). Further, neutrophils display polarization responses to C3A in co-preparations with eosinophils but not when alone, suggesting that C3A stimulates neutrophils indirectly (75). We therefore questioned whether C3A enhances neutrophil migration by sensitizing neutrophils to migrate toward IL-8, a major bioactive neutrophil chemoattractant (76), both in zebrafish wounds(50, 77) and in human wounds and skin graft sites (78, 79). Using microfluidic devices, we found that the total number of neutrophils arriving at the source of an established gradient of IL-8 did not differ between C3A-pre-treated neutrophils and untreated controls, although there was considerable variation between both technical and biological replicates (Fig. 5C). We therefore focused specifically on the neutrophils in each device that eventually reached the chemoattractant source and measured the time at which each neutrophil arrived at the source. Analysis of either the first 20% or first 50% of neutrophils to arrive at the source revealed that neutrophils pre-treated with C3A arrived faster than control neutrophils (Fig 5D). These findings suggest that C3a may prime neutrophils to respond more quickly to other exogenous cues.

### Loss of *c3a.1* impairs neutrophil recruitment *in vivo* by decreasing neutrophil migration speed early after wounding

Our data thus far suggest a role for C3A in priming neutrophils to migrate effectively toward other chemotactic signals. Thus, we next tested whether the impaired neutrophil recruitment phenotype we observed in *c3a.1*^*−/−*^ zebrafish larvae was due to alterations in the dynamics of interstitial migration to the wound. We took advantage of the amenability of larval zebrafish to live time-lapse imaging and single-cell tracking to determine how the interstitial migration characteristics of neutrophils in *c3a.1*-deficient larvae differ from those of wild-type controls. To do this, we crossed the *c3a.1*-deficient line to the *Tg(mpx:mcherry)* line (56), in which neutrophils express the fluorescent protein mcherry. Following tail transection, we imaged labeled neutrophils in the wound microenvironment for 1 hour and tracked the neutrophils using Imaris software (Bitplane) (Fig. 6A, **still images, and movie 2**). We found that average neutrophil speed during the first 30 minutes after wounding is significantly impaired in *c3a.1*^*−/−*^ larvae, compared to *c3a.1*^*+/+*^ controls. This change in neutrophil migratory behavior is confined to the early post-wounding period, as when speed is averaged over the first 60 minutes post-wound, it is not different between groups (Fig. 6B). The mean displacement and track straightness traveled by the neutrophils also did not differ between groups (**Fig. S5A-B**). Decreased neutrophil speed in *c3a.1*^*−/−*^ larvae is specific to neutrophil directed migration, as neutrophil random migration in the absence of an inflammatory stimulus is not impaired in *c3a.1*^*−/−*^ larvae, and neutrophil random migration speed is in fact slightly faster in *c3a.1-*deficient zebrafish than in *c3a.1*^+/+^ controls (**Fig. S5C**). Finally, quantification of each neutrophil’s instantaneous speed at 3 minute intervals over the first hour post-wounding shows that neutrophils in *c3a.1*-intact larvae rapidly achieve and maintain a steady speed toward the wound. In contrast, neutrophils in *c3a.1*^*−/−*^ initially migrate significantly more slowly, before accelerating to reach *c3a.1*^*+/+*^ speeds by about 30 minutes post-wound (Fig. 6C-D). Altogether, these data support the idea that C3A primes neutrophil responses to damaged tissues.

**FIGURE 6.**
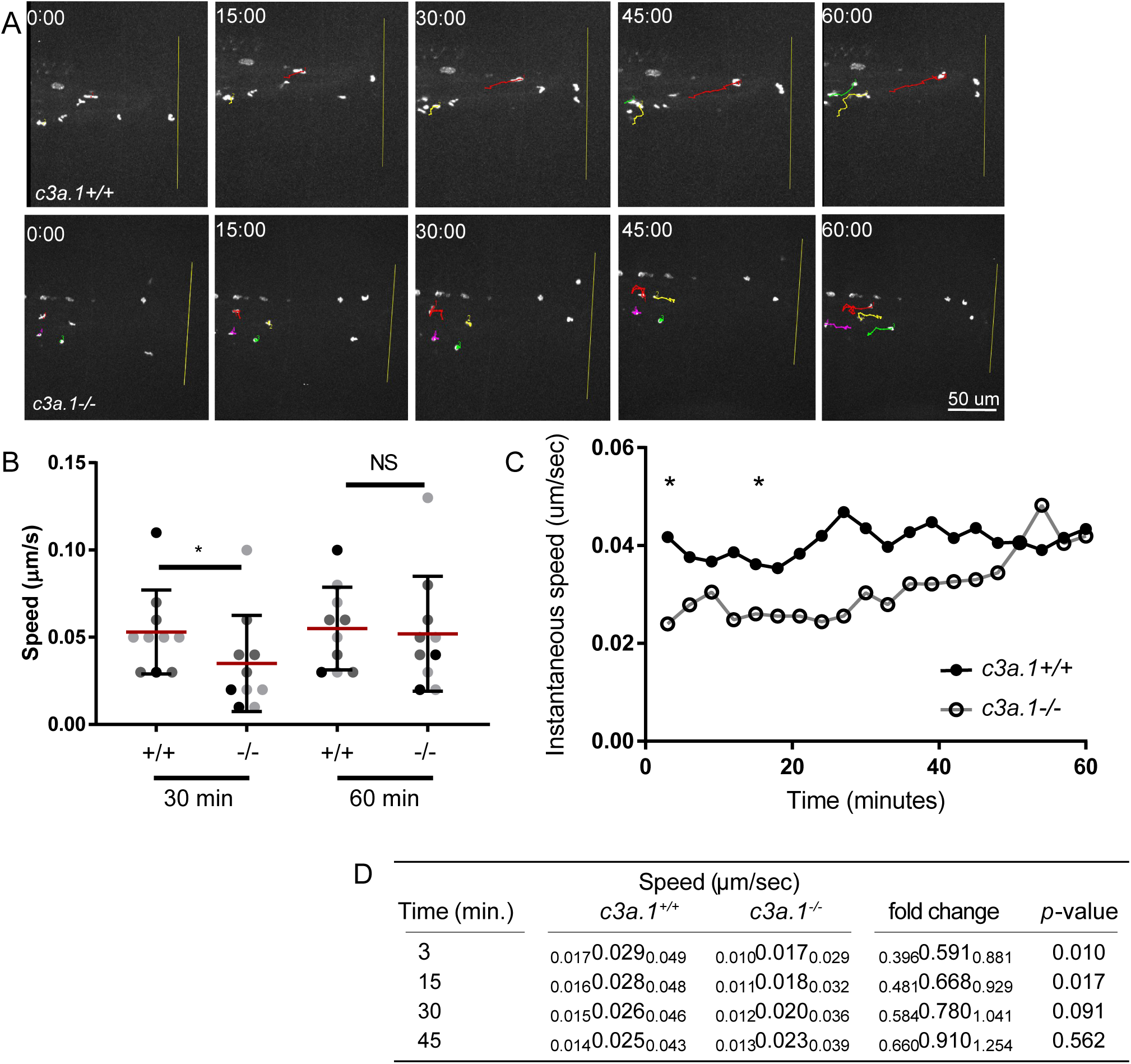
Loss of *c3a.1* impairs neutrophil recruitment by decreasing neutrophil migration speed early after wounding. (A) Time-lapse photomicrographs of neutrophil recruitment to tail-transected caudal fins of *c3a.1*^*+/+*^ (n= 8) or *c3a.1*^*−/−*^ larvae (n = 8) with mcherry-labeled neutrophils (*Tg(mpx:mcherry))*, 0-60 minutes post-wound, showing tracks of forward migrating neutrophils. (B) Quantification of mean track speed of forward-migrating neutrophils, expressed as mean +/− SEM. Each dot represents the mean of the first 5 neutrophils recruited to the wound of an individual larva. Colors represent the results of 4 independent experiments. *p<0.05 (C) Graph, expressed as mean, and (D) quantification of instantaneous speed of all forward-migrating neutrophils over the first 60 minutes following wounding. In (D), for speed and fold change, data are expressed as median (center values), with 95% confidence intervals (small print). Data comprise 4 independent experiments. *p<0.05.

## Discussion

Here we report, for the first time, the results of a cell-specific translation profiling screen designed to identify genes differentially expressed in the inflammatory context of wounding in the larval zebrafish model. We have previously shown that the signals that guide neutrophils to sites of sterile injury differ from those that regulate migration to bacterial infection; specifically, that, while PI3K signaling is required in both contexts, tissue-generated H_2_O_2_ signaling is required for neutrophil responses to wounds, but is dispensable for neutrophil responses to infection (80). Our work here supports the increasing recognition that molecular drivers of innate immune system inflammation are not universal. Context-dependent alterations in the transcriptomes of leukocytes and epithelial cells have the potential to uncover more genes that are differentially expressed only in response to a specific type of inflammatory stimulus. We have also demonstrated the utility of large-scale translation profiling screens in zebrafish to identify promising genes for further study.

We were surprised to find relatively little overlap in the transcriptomes of neutrophils, macrophages, and epithelial cells in that few genes identified by our screen were differentially expressed in more than one cell type. This suggests the presence of a complex network of inter- and intracellular signals, in which cross-communication among cell types is essential for optimal leukocyte recruitment and subsequent wound healing.

We identified *c3a.1* as the only gene significantly upregulated in neutrophils in response to wounding. This finding is interesting because, while the complement system has been implicated in multiple inflammatory contexts, including wounding, infection, and hematopoiesis, it is best understood in infection, where it functions to opsonize bacteria for phagocytosis or kill them directly via assembly of the membrane attack complex (19, 63). Similar to our finding that *c3a.1*^*−/−*^ zebrafish have impaired survival to bacterial infection, mice deficient in either C3 or the C3A receptor have increased susceptibility to septic arthritis (81), and mice with C3A over-activation induced by deletion of the scavenger carboxypeptidase B2 displayed a survival benefit in the context of polymicrobial sepsis (82), confirming a specific, protective role for C3A in infectious inflammatory contexts. However, we find that, in larval zebrafish, C3A acts through neutrophils, as C3A mutation had no further effect when neutrophils were defective.

The role of C3A in the context of sterile injury is less well understood. C3A signaling through the C3A receptor (C3AR) is required for hepatocyte proliferation and liver regeneration following toxic liver injury in mice (83), and C3A can be detected in the wound microenvironment of incised skin wounds in guinea pigs (84). Rafail et al. showed in 2015 that C3^−/−^ mice have faster early wound healing and decreased neutrophil recruitment at wounds than their wild-type counterparts (85). However, while this work showed evidence of a role for the complement pathway in wound-associated inflammation and wound healing, the findings were attributed to a lack of downstream C5a-C5aR1 signaling rather than specific loss of C3A activation (85). Our findings suggest a specific role for C3A in recruiting neutrophils to wounds. We further show that C3A exerts its effects by enhancing neutrophil responses to other chemoattractants, including IL-8.

Because zebrafish C3A and C3B are believed to be the products of different genes and in our *c3a.1*^*−/−*^ model C5A expression is not decreased, our data suggest a specific requirement for C3A in efficient neutrophil recruitment to wounds. However, the length of the zebrafish C3A.1 amino acid sequence is longer than that of human C3A (1643 amino acids in the zebrafish (38), versus 77 amino acids in humans (86)) and contains a consensus sequence for a thioester bond similar to the one cleaved in human C3 to produce C3A and C3B. This suggests that additional, post-translational cleavage occurs in zebrafish to activate C3A.1; alternatively, cleavage of C3A.1 may contribute at least some C3B and downstream complement component activity in zebrafish. It is a limitation of our study that, similar to other authors (48) and most likely due to the large size of the gene, we were not able to express C3A and rescue the neutrophil recruitment phenotype in our *c3a.1*^*−/−*^ model. However, our findings are consistent with the neutrophil recruitment phenotype reported by Forn-Cuni, et al., using *c3a.1* depletion by morpholino (48). Further, in our *in vitro* data generated using human primary neutrophils, we demonstrate that the addition of exogenous C3A induces increased neutrophil chemotaxis to IL-8, a phenotype that is consistent with what we observed *in vivo* with *c3a.1* depletion. Taken together, these findings suggest that C3A plays a role in priming neutrophils for efficient responses to tissue damage cues.

Finally, our data raise additional questions about a neutrophil-specific role for C3A. Our data using *rac2*^*D57N*^ zebrafish mutants suggest that, during infection, impaired survival in *c3a.1*^*−/−*^ larvae is a neutrophil-dependent phenotype. Experiments using either global depletion of *c3a.1* or addition of exogenous C3A show a role for C3A in sensitizing neutrophils to respond to other chemoattractants, such as IL-8; however, *c3a.1* is produced by many cell types (48), and in mammals, C3 is found pre-formed in serum and C3A activated upon injury or infection (19, 73). While our experiments show general effects of C3A on neutrophils, the role of *c3a.1* produced specifically by neutrophils, as indicated by our TRAP-RNAseq results, remains an open question. Autocrine C3AR1 signaling has been implicated in B cell activation and class-switch recombination (87). Neutrophils, as well as other myeloid cells and non-myeloid cells, express C3AR (73, 88), which may be contained in secretory granules and mobilized to the cell surface upon activation (89). Further, neutrophils are able to trigger alternative pathway activation of plasma complement, leading to enhanced neutrophil CD11b expression and respiratory burst (90). However, a specific role for C3AR signaling, autocrine or otherwise, in neutrophil directed migration has not yet been addressed.

In summary, our data identify the complement pathway as a whole, and *c3a.1* in particular, as significantly upregulated in neutrophils in response to wounding. Our data further support a zebrafish model with conserved C3A activity. We demonstrate a role for C3A in priming neutrophils for efficient migration to other chemoattractants, both in vivo in zebrafish and in vitro in human primary neutrophils; however, the role of autocrine neutrophil C3A-C3AR signaling warrants further investigation. On the basis of these observations, we conclude that C3A plays an underappreciated role in mediating neutrophil migration. By exploiting the genetic resources of the larval zebrafish model, we and others are well positioned to further investigate the role of neutrophil-derived C3A in optimizing neutrophil recruitment to wounding. Finally, these results support the power of TRAP in the identification of cell type specific changes in gene expression that may influence inflammation and wound healing.

## Supplemental Data

**Table 1:** Published zebrafish lines used in this study.

**Table 2:** Primer sequences used in this study.

**Table 3:** Differentially expressed genes shown in Figure 1 C-D.

**S1:** Global depletion of *c3a.1* delays tail development and decreases regenerate length following tail transection. Quantification of caudal fin length during larval development, 4dpf-6dpf, of *c3a.1*^*+/+*^ (n = 40, 4dpf; 40, 5dpf; 19, 6dpf) and *c3a.1*^*−/−*^ (n = 36, 4dpf; 34, 5dpf; 13, 6dpf) larvae. All data are expressed as mean +/− SEM, with each dot representing 1 larva and colors representing the results of 3 independent experiments. **p<0.01.

**S2.** Global depletion of *c3a.1* does not alter macrophage recruitment to tail wounds. Quantification of macrophage numbers at the wound following tail transection of *c3a.1*^*+/+*^ (n = 15, 4hpw; 10, 24hpw) or *c3a.1*^*−/−*^ (n = 19, 4hpw; 17, 24hpw) *Tg(mpeg1:GFP)* larvae, expressed as mean +/−SEM. Each dot represents one larva; colors represent the results of 3 independent experiments.

**S3.** Fin regeneration is impaired after tail transection in *c3a.1*^*−/−*^ larvae. Quantification of caudal fin regenerate length following tail transection, 24 hpw-72 hpw, of *c3a.1*^*+/+*^ (n = 20, 24 hpw; 20, 48 hpw; 36, 72 hpw) and *c3a.1*^*−/−*^ (n = 39, 24 hpw; 40, 48 hpw; 36, 72 hpw) larvae. All data are expressed as mean +/− SEM, with each dot representing 1 larva and colors representing the results of 3 independent experiments. **p<0.01, ****p<0.0001.

**S4.** (A) CHT neutrophils of *Tg(mpx:dendra2) c3a.1*^*+/+*^ or *c3a.1*^*−/−*^ were photoconverted and the larvae subjected to tail transection. (B) Quantification of photoconverted neutrophils retained in the CHT at 3hpw. (C) Quantification of photoconverted neutrophils outside the CHT at 3hpw. *c3a.1*^*+/+*^ (n=16); *c3a.1*^*−/−*^ (n=22). All data are expressed as mean +/− SEM, with each dot representing one larva and colors representing 4 independent experiments.

**Movie 1.** (A) Representative time-lapse movies of human primary neutrophils exposed to a gradient of either C5a (3 uM source, left) or C3a (3 uM source, right). The source of the gradient is located center-bottom. (B) Representative time-lapse movies of human primary neutrophils exposed to a gradient of IL-8 (1 uM source) following pretreatment with either media (left) or 3 uM C3a (right). The source gradient is located center-bottom.

**Movie 2.** Representative time-lapse movies of recruitment of mcherry-labeled neutrophils to tail-transected caudal fins of *c3a.1*^*+/+*^ (left) or *c3a.1*^*−/−*^ (right) zebrafish larvae, 0-60 minutes post-wound. Tracks of forward-migrating neutrophils are labeled in color.

**S5.** (A) Quantification of mean linear displacement distance traveled by forward-migrating neutrophils in *c3a.1*^*+/+*^ (n = 8) and *c3a.1*^*−/−*^ (n = 8) during early neutrophil recruitment. (B) Mean track straightness of forward-migrating neutrophils during early neutrophil recruitment does not differ between *c3a.1*^*+/+*^ (n = 8) and *c3a.1*^*−/−*^ (n = 8) larvae. (C) Quantification of mean track speed of randomly migrating neutrophils in the heads of unwounded *c3a.1*^*+/+*^ (n = 9) or *c3a.1*^*−/−*^ (n = 8) larvae. All data are expressed as mean +/− SEM. Each dot represents the mean of value for all neutrophils for one larva. Colors represent results of 4 independent experiments. *p<0.05.

## Acknowledgements

We would like to thank Christina M. Freisinger and Elizabeth A. Harvie for assistance with the TRAP sample preparation. We would also like to acknowledge the UW-Madison Biotechnology Center for assistance with library preparation and Illumina sequencing. We thank Michael R. Lasarev from the UW-Madison Department of Biostatistics and Medical Informatics for additional support provided through the Clinical and Translation Science Award (CTSA) program, grant UL1TR002373. This work was funded by NIH R35 GM1 18027 01 (AH). RAH was supported by an individual fellowship under NIH T32HL07899.

