## Supplementary figures and images for "Cell type specific gene expression profiling reveals a role for the complement component C3A in neutrophil migration to tissue damage"

### Supplemental Figure 1

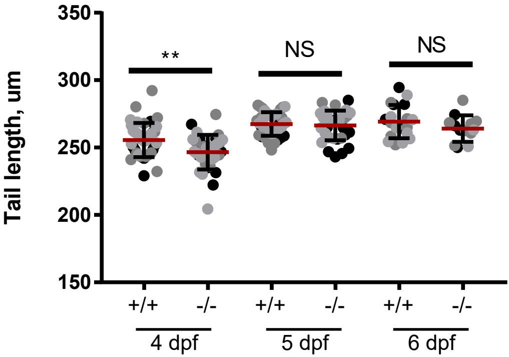

### Supplemental figure 2

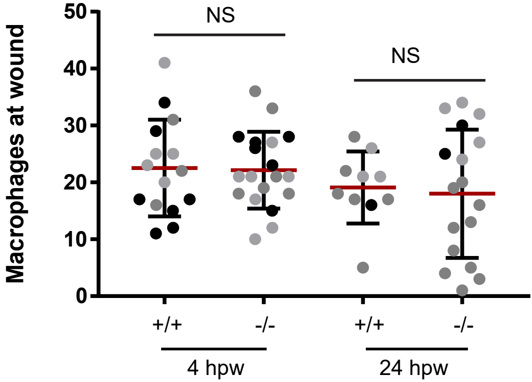

### Supplemental figure 3

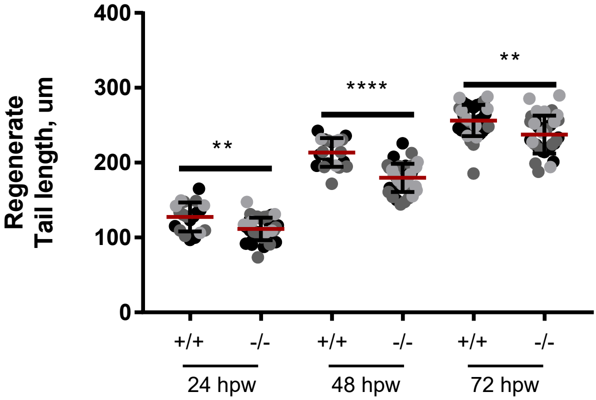

### Supplemental figure 4

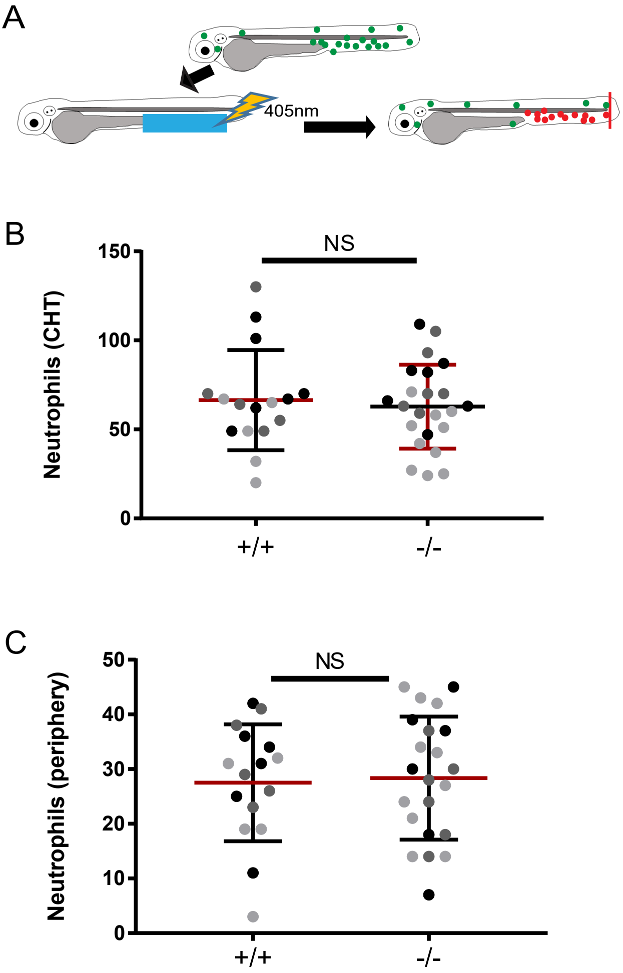

### Supplemental figure 5

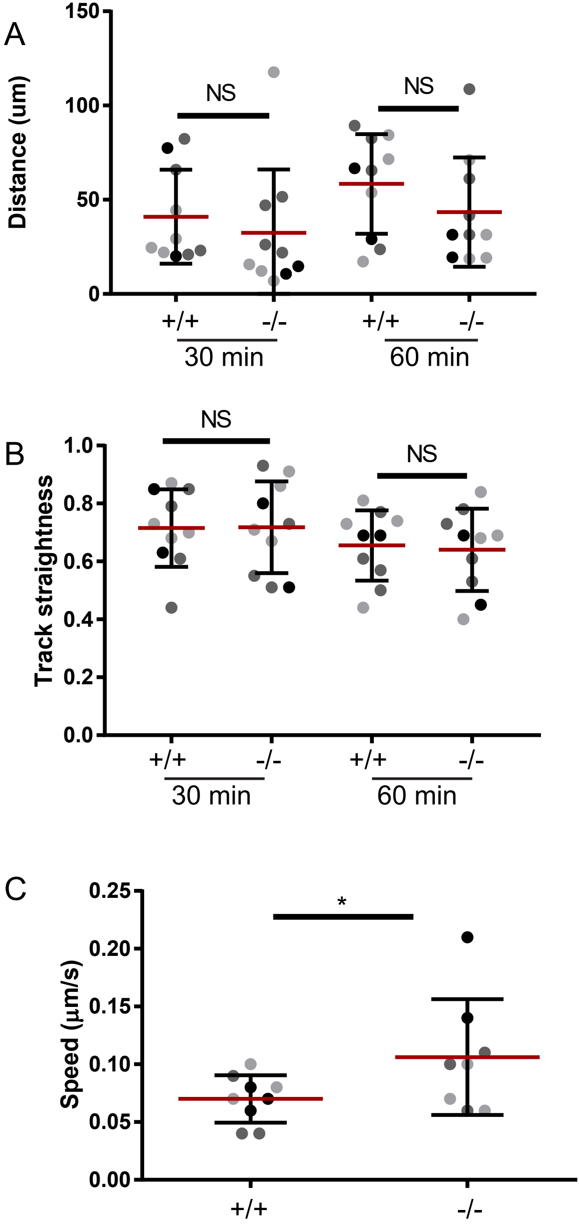
