## Supplemental Table 1 for "Cell type specific gene expression profiling reveals a role for the complement component C3A in neutrophil migration to tissue damage"

| Line | Reference |
|---|---|
| lyz:L10a-EGFP | (17) |
| mpeg1:L10a-EGFP | (17) |
| krt4:L10a-EGFP | (17) |
| mpeg1:EGFP | (13) |
| mpx:mCherry-2A-rac2 | (51) |
| mpx:mCherry-2A-rac2$^{D57N}$ | (51) |
| mpx:Dendra2 | (54) |
| mpx:mCherry | (56) |
