## Supplemental table 2 for "Cell type specific gene expression profiling reveals a role for the complement component C3A in neutrophil migration to tissue damage"

| Primer | Sequence (5' to 3') | Ref. (if prev. published) |
|---|---|---|
| sa31241_F | TCACTCACGCTCTGTCTCTC | |
| sa31241_R | GGAAACATAGCTACTGACTGGA | |
| C3.1 qPCR pub F | TCCAGACAAGCGAAAGGTG | (48) |
| C3.1 qPCR pub R | CCATCAGTGTACACAGCATCATAC | (48) |
| C3.2/3 qPCR F | CGGTACACAAACACCCCTCT | (48) |
| C3.2/3 qPCR R | GTCTTCCTCATCGTTCTCTTGTT | (48) |
| C3.4 qPCR F | CAACTCAGAAGCGTCCATGA | (48) |
| C3.4 qPCR R | ATTGATCAGCCCTTGCAACT | (48) |
| C3.5 qPCR F | GTTGCACGCACAGACAAGTT | (48) |
| C3.5 qPCR R | CAGGCTCTTTCTCCATCTGC | (48) |
| C3.6 qPCR F | CAGACCACATCACTGCCAAC | (48) |
| C3.6 qPCR R | TTGTGCATCCGAAGTTGAAG | (48) |
| C3.7/8 qPCR F | CTCCATTTCGATGGCTGAAT | (48) |
| C3.7/8 qPCR R | ACATCACTCCGACCAGGAAC | (48) |
| c3b.1 qPCR F | TGAGATGGAGATTGTGCAGGT | |
| c3b.1 qPCR R | CTGCAGCTTGCATGAGAGAG | |
| c3b.2 qPCR F | CTGATCAGCATCAGCCAGAG | |
| c3b.2 qPCR R | CGCATGAGAGAGGAACATCC | |
| C5 qPCR F | CGGTTCAATCAGTGCTCAAA | |
| C5 qPCR R | TACTGCTTGCCAATCTCGAA | |
| EF1a qPCR F | TGCCTTCGTCCCAATTTCAG | (49) |
| EF1a qPCR R | TACCCTCCTTGCGCTCAATC | (49) |
