## Supplemental table 3A for "Cell type specific gene expression profiling reveals a role for the complement component C3A in neutrophil migration to tissue damage"

| Abs fold change > 2 after wounding (only genes with human orthologs) | | | | | | | | |
|---|---|---|---|---|---|---|---|---|
| Neutrophils (lyz) | | | Macrophages (mpeg1) | | | Epithelium (krt4) | | |
| **Ensembl gene ID** | **Gene name** | **Human ortholog ID** | **Ensembl gene ID** | **Gene name** | **Human ortholog ID** | **Ensembl gene ID** | **Gene name** | **Human ortholog ID** |
| ENSDARG00000016319 | c9 | ENSG00000113600 | ENSDARG00000101762 | fah | ENSG00000103876 | ENSDARG00000057273 | alox5a | ENSG00000012779 |
| ENSDARG00000021004 | c5 | ENSG00000106804 | ENSDARG00000101530 | prr16 | ENSG00000184838 | ENSDARG00000100376 | amdhd2 | ENSG00000162066 |
| ENSDARG00000070723 | ndufa9a | ENSG00000139180 | ENSDARG00000053646 | adrb3a | ENSG00000188778 | ENSDARG00000062101 | iffo2a | ENSG00000169991 |
| ENSDARG00000103370 | lrrc8da | ENSG00000171017 | ENSDARG00000019001 | ak7a | ENSG00000140057 | ENSDARG00000031119 | baiap2l1b | ENSG00000006453 |
| ENSDARG00000012694 | c3a.1 | ENSG00000125730 | ENSDARG00000101557 | scarb1 | ENSG00000073060 | ENSDARG00000030263 | mfsd2b | ENSG00000205639 |
| ENSDARG00000051896 | fbn2a | ENSG00000138829 | ENSDARG00000075067 | aipl1 | ENSG00000129221 | ENSDARG00000010312 | cp | ENSG00000047457 |
| ENSDARG00000077261 | lmod1b | ENSG00000163431 | ENSDARG00000023886 | cacna2d4b | ENSG00000151062 | ENSDARG00000088672 | CABZ01054962.1 | ENSG00000089692 |
| ENSDARG00000036457 | cacng6a | ENSG00000130433 | ENSDARG00000032831 | htra1a | ENSG00000166033 | ENSDARG00000097630 | C24H8orf59 | ENSG00000176731 |
| ENSDARG00000007494 | st3gal8 | ENSG00000125975 | ENSDARG00000069844 | irg1 | ENSG00000102794 | ENSDARG00000098904 | pex1 | ENSG00000127980 |
| ENSDARG00000031119 | baiap2l1b | ENSG00000006453 | ENSDARG00000004618 | stx2a | ENSG00000111450 | ENSDARG00000054032 | rab24 | ENSG00000169228 |
| ENSDARG00000067672 | card9 | ENSG00000187796 | ENSDARG00000053003 | slc35f1 | ENSG00000196376 | ENSDARG00000077569 | lrrn3 | ENSG00000173114 |
| ENSDARG00000076950 | kel | ENSG00000196549 | ENSDARG00000044895 | fmoda | ENSG00000122176 | ENSDARG00000045706 | tspan9b | ENSG00000011105 |
| ENSDARG00000058180 | exoc3l4 | ENSG00000205436 | ENSDARG00000055514 | icn2 | ENSG00000188643 | ENSDARG00000074254 | abcb6b | ENSG00000115657 |
| ENSDARG00000057276 | iqca1 | ENSG00000132321 | ENSDARG00000055970 | zgc:136971 | ENSG00000197635 | ENSDARG00000061031 | dner | ENSG00000187957 |
| ENSDARG00000102332 | SPINT1 (2 of 2) | ENSG00000101448 | ENSDARG00000043177 | lsm1 | ENSG00000175324 | ENSDARG00000052035 | nxnl1 | ENSG00000171773 |
| ENSDARG00000044433 | sned1 | ENSG00000162804 | ENSDARG00000006981 | ccdc113 | ENSG00000103021 | ENSDARG00000077009 | wdfy4 | ENSG00000128815 |
| ENSDARG00000071065 | g6pd | ENSG00000160211 | ENSDARG00000022399 | ntmt1 | ENSG00000148335 | ENSDARG00000104801 | tubb6 | ENSG00000176014 |
| ENSDARG00000019950 | carf | ENSG00000138380 | ENSDARG00000070918 | FETUB (2 of 2) | ENSG00000090512 | ENSDARG00000074428 | hhat | ENSG00000054392 |
| ENSDARG00000055278 | cfb | ENSG00000243649 | ENSDARG00000023645 | ccbl1 | ENSG00000171097 | ENSDARG00000010918 | nprl3 | ENSG00000103148 |
| ENSDARG00000054150 | cx23 | ENSG00000203733 | ENSDARG00000040466 | vil1 | ENSG00000127831 | ENSDARG00000056582 | MAP3K14 (1 of 2) | ENSG00000006062 |
| ENSDARG00000091349 | plekhd1 | ENSG00000175985 | ENSDARG00000038397 | rpap2 | ENSG00000122484 | ENSDARG00000097795 | si:ch73-195i19.3 | ENSG00000169255 |
| ENSDARG00000101242 | zcchc14 | ENSG00000140948 | ENSDARG00000045594 | si:dkey-14k9.2 | ENSG00000138944 | ENSDARG00000026224 | fanci | ENSG00000140525 |
| ENSDARG00000019951 | sec62 | ENSG00000008952 | ENSDARG00000015293 | fam110a | ENSG00000125898 | ENSDARG00000003411 | foxa2 | ENSG00000125798 |
| ENSDARG00000011449 | fes | ENSG00000182511 | ENSDARG00000005799 | smim14 | ENSG00000163683 | ENSDARG00000014976 | lims2 | ENSG00000169756 |
| ENSDARG00000011506 | lpcat1 | ENSG00000153395 | ENSDARG00000014320 | gucy2c | ENSG00000070019 | ENSDARG00000069844 | irg1 | ENSG00000102794 |
| ENSDARG00000078941 | wu:fe11b02 | ENSG00000164678 | ENSDARG00000104259 | RAB26 | ENSG00000167964 | ENSDARG00000040036 | tcf19l | ENSG00000137310 |
| ENSDARG00000056617 | rpgra | ENSG00000156313 | ENSDARG00000032631 | ltb4r | ENSG00000213903 | ENSDARG00000017007 | nr4a2a | ENSG00000153234 |
| ENSDARG00000090833 | igfbp6b | ENSG00000167779 | ENSDARG00000052129 | boka | ENSG00000176720 | ENSDARG00000004445 | GRM5 (1 of 2) | ENSG00000168959 |
| ENSDARG00000042833 | rffl | ENSG00000092871 | ENSDARG00000104868 | KCNT1 (2 of 2) | ENSG00000107147 | ENSDARG00000062959 | | ENSG00000173926 |
| ENSDARG00000023026 | pkp2 | ENSG00000057294 | ENSDARG00000076376 | tmem175 | ENSG00000127419 | ENSDARG00000074904 | DBNDD2 | ENSG00000244274 |
| ENSDARG00000031136 | moxd1 | ENSG00000079931 | ENSDARG00000008127 | pcdh15b | ENSG00000150275 | ENSDARG00000102302 | WDR48 (2 of 3) | ENSG00000114742 |
| ENSDARG00000042630 | hebp2 | ENSG00000051620 | ENSDARG00000052264 | chrnb3a | ENSG00000147432 | ENSDARG00000098761 | rgs2 | ENSG00000116741 |
| ENSDARG00000032126 | scg5 | ENSG00000166922 | ENSDARG00000090625 | BCL2L12 | ENSG00000126453 | ENSDARG00000077654 | grm8a | ENSG00000179603 |
| ENSDARG00000102783 | slc25a35 | ENSG00000125434 | ENSDARG00000054150 | cx23 | ENSG00000203733 | ENSDARG00000103390 | CACNA2D2 (3 of | ENSG00000007402 |
| ENSDARG00000028628 | gdap1l1 | ENSG00000124194 | ENSDARG00000067613 | dnajc25 | ENSG00000059769 | ENSDARG00000053475 | ngb | ENSG00000165553 |
| ENSDARG00000070100 | RIN1 (1 of 2) | ENSG00000174791 | ENSDARG00000039827 | bbs5 | ENSG00000163093 | ENSDARG00000056938 | kera | ENSG00000139330 |
| ENSDARG00000039650 | itm2cb | ENSG00000135916 | ENSDARG00000071573 | lsm5 | ENSG00000106355 | ENSDARG00000032780 | pigf | ENSG00000151665 |
| ENSDARG00000068516 | hapln1b | ENSG00000145681 | ENSDARG00000101024 | nufip1 | ENSG00000083635 | ENSDARG00000068286 | slc7a4 | ENSG00000099960 |
| ENSDARG00000094473 | FHAD1 | ENSG00000142621 | ENSDARG00000057169 | abca4a | ENSG00000198691 | ENSDARG00000090526 | PLEKHS1 (3 of 3) | ENSG00000148735 |
| ENSDARG00000067797 | spi1a | ENSG00000142539 | ENSDARG00000075833 | lyve1a | ENSG00000133800 | ENSDARG00000058042 | kitlgb | ENSG00000049130 |
| ENSDARG00000098311 | ZFOS-932H1.3 | ENSG00000169740 | ENSDARG00000100381 | rock2bl | ENSG00000134318 | ENSDARG00000068296 | SLCO3A1 (2 of 2) | ENSG00000176463 |
| ENSDARG00000002945 | bgnb | ENSG00000182492 | ENSDARG00000103647 | gpc5a | ENSG00000179399 | ENSDARG00000059523 | rims4 | ENSG00000101098 |
| ENSDARG00000088950 | sycp2 | ENSG00000196074 | ENSDARG00000098700 | il1b | ENSG00000125538 | ENSDARG00000027957 | fgf12a | ENSG00000114279 |
| ENSDARG00000006397 | laptm4a | ENSG00000068697 | ENSDARG00000014973 | ntng1a | ENSG00000162631 | ENSDARG00000030215 | matn1 | ENSG00000162510 |
| ENSDARG00000101989 | THRAP3 (2 of 2) | ENSG00000054118 | ENSDARG00000075008 | pask | ENSG00000115687 | ENSDARG00000056950 | epyc | ENSG00000083782 |
| ENSDARG00000017953 | tp73 | ENSG00000078900 | ENSDARG00000044815 | SERINC4 | ENSG00000184716 | ENSDARG00000078989 | alpk3a | ENSG00000136383 |
| ENSDARG00000061690 | nxpe3 | ENSG00000144815 | ENSDARG00000099840 | lpar1 | ENSG00000198121 | ENSDARG00000022176 | IGLON5 | ENSG00000142549 |
| ENSDARG00000077202 | mocs2 | ENSG00000164172 | ENSDARG00000026387 | tmem55bb | ENSG00000165782 | ENSDARG00000052025 | pomgnt1 | ENSG00000085998 |
| ENSDARG00000062987 | tyw1 | ENSG00000198874 | ENSDARG00000071203 | sptssa | ENSG00000165389 | ENSDARG00000060861 | dars2 | ENSG00000117593 |
| ENSDARG00000053520 | C12H7orf62 | ENSG00000164645 | ENSDARG00000078989 | alpk3a | ENSG00000136383 | ENSDARG00000077863 | mettl22 | ENSG00000067365 |
| ENSDARG00000088877 | smarcd2 | ENSG00000108604 | ENSDARG00000058318 | c1qtnf9 | ENSG00000205863 | ENSDARG00000076082 | CABZ01041962.1 | ENSG00000173451 |
| ENSDARG00000078508 | KCP | ENSG00000135253 | ENSDARG00000074892 | pde6d | ENSG00000156973 | ENSDARG00000008757 | entpd6 | ENSG00000197586 |
| ENSDARG00000057859 | hoxd10a | ENSG00000128710 | ENSDARG00000087517 | btbd3b | ENSG00000132640 | ENSDARG00000051896 | fbn2a | ENSG00000138829 |
| ENSDARG00000077139 | col6a3 | ENSG00000163359 | ENSDARG00000055416 | serpinb1 | ENSG00000021355 | ENSDARG00000089844 | SCARB2 | ENSG00000138760 |
| ENSDARG00000079463 | MAN2B2 | ENSG00000013288 | ENSDARG00000069009 | si:ch211-147a11.: | ENSG00000198912 | ENSDARG00000036186 | mbpa | ENSG00000197971 |
| ENSDARG00000091260 | mylk4a | ENSG00000145949 | ENSDARG00000055252 | snap23.2 | ENSG00000092531 | ENSDARG00000029865 | rassf2a | ENSG00000101265 |
| ENSDARG00000078094 | lmf2a | ENSG00000100258 | ENSDARG00000014910 | panx1b | ENSG00000110218 | ENSDARG00000056651 | gfra4a | ENSG00000125861 |
| ENSDARG00000036767 | si:dkey-204a24.1( | ENSG00000106608 | ENSDARG00000100157 | tamm41 | ENSG00000144559 | ENSDARG00000039852 | ftr93 | ENSG00000128250 |
| ENSDARG00000011410 | AADAT | ENSG00000109576 | ENSDARG00000055070 | BORCS7 | ENSG00000166275 | ENSDARG00000007366 | amdhd1 | ENSG00000139344 |
| ENSDARG00000043055 | hiat1b | ENSG00000156875 | ENSDARG00000058292 | sephs1 | ENSG00000086475 | ENSDARG00000003054 | tacr3l | ENSG00000169836 |
| ENSDARG00000100203 | SZT2 | ENSG00000198198 | ENSDARG00000028367 | sult2st3 | ENSG00000088002 | ENSDARG00000069196 | ebf1b | ENSG00000164330 |
| ENSDARG00000102302 | WDR48 (2 of 3) | ENSG00000114742 | ENSDARG00000088672 | CABZ01054962.1 | ENSG00000089692 | ENSDARG00000007009 | hoxa11b | ENSG00000005073 |
| ENSDARG00000100304 | chdh | ENSG00000016391 | ENSDARG00000040350 | rpp38 | ENSG00000152464 | ENSDARG00000078163 | hpse2 | ENSG00000172987 |
| ENSDARG00000029688 | hsp70.1 | ENSG00000126803 | ENSDARG00000039699 | fam204a | ENSG00000165669 | ENSDARG00000074283 | inpp5d | ENSG00000168918 |
| ENSDARG00000003219 | bin2a | ENSG00000110934 | ENSDARG00000025034 | syngr3b | ENSG00000127561 | ENSDARG00000101979 | eps8l3a | ENSG00000198758 |
| ENSDARG00000029036 | rab32a | ENSG00000118508 | ENSDARG00000077606 | arsia | ENSG00000183876 | ENSDARG00000079024 | rps6kl1 | ENSG00000198208 |
| ENSDARG00000056450 | MANEAL | ENSG00000185090 | ENSDARG00000101363 | ano2 | ENSG00000047617 | ENSDARG00000052527 | mul1b | ENSG00000090432 |
| ENSDARG00000041394 | dnajb1b | ENSG00000132002 | ENSDARG00000012460 | rassf4 | ENSG00000107551 | ENSDARG00000078060 | syt7b | ENSG00000011347 |
| ENSDARG00000008969 | fgb | ENSG00000171564 | ENSDARG00000022303 | higd1a | ENSG00000131097 | ENSDARG00000041530 | ppih | ENSG00000171960 |
| ENSDARG00000076790 | crygm2d16 | ENSG00000118231 | ENSDARG00000101061 | eif2a | ENSG00000144895 | ENSDARG00000069698 | lrguk | ENSG00000155530 |
| ENSDARG00000103774 | limch1b | ENSG00000064042 | ENSDARG00000087301 | crygm2d14 | ENSG00000118231 | ENSDARG00000103457 | spg20b | ENSG00000133104 |
| ENSDARG00000040274 | scamp5b | ENSG00000198794 | ENSDARG00000090106 | slc17a7b | ENSG00000104888 | ENSDARG00000033999 | cyp26a1 | ENSG00000095596 |
| ENSDARG00000020741 | fga | ENSG00000171560 | ENSDARG00000100231 | ccdc135 | ENSG00000159625 | ENSDARG00000071087 | slc35b4 | ENSG00000205060 |
| ENSDARG00000042291 | dlx6a | ENSG00000006377 | ENSDARG00000031422 | igfbp2b | ENSG00000115457 | ENSDARG00000028276 | | ENSG00000253148 |
| ENSDARG00000031044 | lipg | ENSG00000101670 | ENSDARG00000094377 | samd10b | ENSG00000130590 | ENSDARG00000030020 | agfg1a | ENSG00000173744 |
| ENSDARG00000063525 | adpgk | ENSG00000159322 | ENSDARG00000060668 | tm4sf18 | ENSG00000163762 | ENSDARG00000078094 | lmf2a | ENSG00000100258 |
| ENSDARG00000030933 | ksr1b | ENSG00000141068 | ENSDARG00000075726 | dhrs7b | ENSG00000109016 | ENSDARG00000036755 | prmt9 | ENSG00000164169 |
| ENSDARG00000042180 | foxl2 | ENSG00000183770 | ENSDARG00000104519 | KDELR3 (2 of 2) | ENSG00000100196 | ENSDARG00000061222 | uba7 | ENSG00000130985 |
| ENSDARG00000052644 | ca10a | ENSG00000154975 | ENSDARG00000030012 | lrrfip1a | ENSG00000124831 | ENSDARG00000029609 | macrod1 | ENSG00000133315 |
| ENSDARG00000008433 | unc45b | ENSG00000141161 | ENSDARG00000022378 | mtmr1b | ENSG00000063601 | ENSDARG00000037498 | gria3b | ENSG00000125675 |
| ENSDARG00000038788 | dnai1.2 | ENSG00000122735 | ENSDARG00000053934 | hdhd3 | ENSG00000119431 | ENSDARG00000062633 | cadm2b | ENSG00000175161 |
| ENSDARG00000032780 | pigf | ENSG00000151665 | ENSDARG00000099425 | CFI | ENSG00000205403 | ENSDARG00000063019 | panx2 | ENSG00000073150 |
| ENSDARG00000041779 | trdn | ENSG00000186439 | ENSDARG00000060435 | bnipl | ENSG00000163141 | ENSDARG00000093359 | inpp5jb | ENSG00000185133 |
| ENSDARG00000010794 | tfeb | ENSG00000112561 | ENSDARG00000086647 | chrng | ENSG00000196811 | ENSDARG00000095949 | si:dkey-22i16.9 | ENSG00000107738 |
| ENSDARG00000102021 | C17H14orf28 | ENSG00000179476 | ENSDARG00000100455 | pnkp | ENSG00000039650 | ENSDARG00000018721 | npnt | ENSG00000168743 |
| ENSDARG00000018621 | slc6a19a.1 | ENSG00000174358 | ENSDARG00000032199 | gpc3 | ENSG00000147257 | ENSDARG00000060152 | fam155b | ENSG00000130054 |
| ENSDARG00000022971 | epha6 | ENSG00000080224 | ENSDARG00000029239 | syt9b | ENSG00000170743 | ENSDARG00000102687 | NBL1 | ENSG00000158747 |
| ENSDARG00000016412 | agt | ENSG00000135744 | ENSDARG00000051737 | zgc:173552 | ENSG00000168148 | ENSDARG00000094901 | ABCC6 | ENSG00000121270 |
| ENSDARG00000070059 | si:ch211-262i1.5 | ENSG00000174796 | ENSDARG00000079124 | MTERF4 | ENSG00000122085 | ENSDARG00000041414 | bmf2 | ENSG00000104081 |
| ENSDARG00000070726 | cnga3a | ENSG00000144191 | ENSDARG00000069843 | kctd12.1 | ENSG00000178695 | ENSDARG00000043796 | ms4a17a.7 | ENSG00000110077 |
| ENSDARG00000037276 | map9 | ENSG00000164114 | ENSDARG00000016044 | gins4 | ENSG00000147536 | ENSDARG00000061899 | iqub | ENSG00000164675 |
| ENSDARG00000018553 | ifih1 | ENSG00000115267 | ENSDARG00000007356 | fgf20a | ENSG00000078579 | ENSDARG00000074765 | mhc1zja | ENSG00000204525 |
| ENSDARG00000027345 | mpzl2b | ENSG00000149573 | ENSDARG00000059053 | slc13a4 | ENSG00000164707 | ENSDARG00000058985 | KCNJ6 | ENSG00000157542 |
| ENSDARG00000092362 | hsp70.2 | ENSG00000126803 | ENSDARG00000001721 | oc90 | ENSG00000253117 | ENSDARG00000019556 | clcn7 | ENSG00000103249 |
| ENSDARG00000055723 | hsp70l | ENSG00000126803 | ENSDARG00000058365 | hspb8 | ENSG00000152137 | ENSDARG00000100897 | PDGFRB | ENSG00000113721 |
| ENSDARG00000001442 | inppl1b | ENSG00000198574 | ENSDARG00000101800 | PSTK | ENSG00000179988 | ENSDARG00000013006 | pibf1 | ENSG00000083535 |
| ENSDARG00000092882 | si:dkey-161j23.7 | ENSG00000047578 | ENSDARG00000078378 | TRAF2 (2 of 2) | ENSG00000127191 | ENSDARG00000038446 | nrp2b | ENSG00000118257 |
| ENSDARG00000041294 | noxo1a | ENSG00000196408 | ENSDARG00000035798 | gngt1 | ENSG00000127920 | ENSDARG00000103826 | gpib | ENSG00000105220 |

| | | |
|---|---|---|
| ENSDARG00000061899 | iqub | ENSG00000164675 |
| ENSDARG00000071087 | slc35b4 | ENSG00000205060 |
| ENSDARG00000062058 | slc12a7b | ENSG00000113504 |
| ENSDARG00000075720 | il2rb | ENSG00000100385 |
| ENSDARG00000069843 | kctd12.1 | ENSG00000178695 |
| ENSDARG00000058042 | kitlgb | ENSG00000049130 |
| ENSDARG00000078052 | emilin3a | ENSG00000183798 |
| ENSDARG00000014003 | stard8 | ENSG00000130052 |
| ENSDARG00000057983 | svopl | ENSG00000157703 |
| ENSDARG00000079796 | ldah | ENSG00000118961 |
| ENSDARG00000077058 | CELA1 (6 of 7) | ENSG00000139610 |
| ENSDARG00000057986 | ints8 | ENSG00000164941 |
| ENSDARG00000052089 | CCKAR | ENSG00000163394 |
| ENSDARG00000075672 | pms2 | ENSG00000122512 |
| ENSDARG00000077982 | elf3 | ENSG00000163435 |
| ENSDARG00000024090 | dnajc4 | ENSG00000110011 |
| ENSDARG00000062081 | tbc1d1 | ENSG00000065882 |
| ENSDARG00000068397 | tns2b | ENSG00000111077 |
| ENSDARG00000103600 | st8sia4 | ENSG00000113532 |
| ENSDARG00000098720 | LPL (2 of 2) | ENSG00000175445 |
| ENSDARG00000073756 | slc12a7a | ENSG00000113504 |
| ENSDARG00000101178 | SH3RF2 (2 of 2) | ENSG00000154447 |
| ENSDARG00000103983 | slc30a10 | ENSG00000196660 |
| ENSDARG00000069669 | adra2c | ENSG00000184160 |
| ENSDARG00000007382 | ubtd1a | ENSG00000165886 |
| ENSDARG00000021287 | zgc:91909 | ENSG00000075785 |
| ENSDARG00000053010 | lpcat2 | ENSG00000087253 |
| ENSDARG00000044094 | gfpt2 | ENSG00000131459 |
| ENSDARG00000100700 | EPS15L1 (2 of 2) | ENSG00000127527 |
| ENSDARG00000035918 | bzw2 | ENSG00000136261 |
| ENSDARG00000013542 | lpgat1 | ENSG00000123684 |
| ENSDARG00000021547 | vrk2 | ENSG00000028116 |
| ENSDARG00000027572 | as3mt | ENSG00000214435 |
| ENSDARG00000040700 | nab1b | ENSG00000138386 |
| ENSDARG00000098021 | si:dkey-111k8.2 | ENSG00000167394 |
| ENSDARG00000038938 | ssfa2 | ENSG00000138434 |
| ENSDARG00000062902 | TULP2 | ENSG00000104804 |
| ENSDARG00000004171 | rabac1 | ENSG00000105404 |
| ENSDARG00000019438 | rnf13 | ENSG00000082996 |
| ENSDARG00000011821 | plod2 | ENSG00000152952 |
| ENSDARG00000063100 | psmd14 | ENSG00000115233 |
| ENSDARG00000014646 | aoc2 | ENSG00000131471 |
| ENSDARG00000020278 | si:dkeyp-120h9.1 | ENSG00000164893 |
| ENSDARG00000013926 | slc16a9a | ENSG00000165449 |
| ENSDARG00000078789 | tmem184c | ENSG00000164168 |
| ENSDARG00000104550 | CLCN2 (3 of 3) | ENSG00000114859 |
| ENSDARG00000060029 | fam160b2 | ENSG00000158863 |
| ENSDARG00000059729 | si:ch211-89f7.1 | ENSG00000100003 |
| ENSDARG00000045463 | ralaa | ENSG00000006451 |
| ENSDARG00000101715 | arsg | ENSG00000141337 |
| ENSDARG00000074867 | STIM2 (2 of 2) | ENSG00000109689 |
| ENSDARG00000028393 | daam2 | ENSG00000146122 |
| ENSDARG00000003217 | urb2 | ENSG00000135763 |
| ENSDARG00000019496 | esd | ENSG00000139684 |
| ENSDARG00000103786 | klhl13 | ENSG00000003096 |
| ENSDARG00000019797 | qrich1 | ENSG00000198218 |
| ENSDARG00000062519 | abcc13 | ENSG00000091262 |
| ENSDARG00000071679 | mydgf | ENSG00000074842 |
| ENSDARG00000035985 | FAM210A (1 of 2) | ENSG00000177150 |
| ENSDARG00000078775 | RASA2 | ENSG00000155903 |
| ENSDARG00000092660 | cyp27c1 | ENSG00000186684 |
| ENSDARG00000071683 | zfr2 | ENSG00000105278 |
| ENSDARG00000052030 | akr1a1b | ENSG00000117448 |
| ENSDARG00000074265 | dzip1l | ENSG00000158163 |
| ENSDARG00000030129 | HIATL1 (2 of 2) | ENSG00000148110 |
| ENSDARG00000041947 | styk1 | ENSG00000060140 |
| ENSDARG00000061974 | grhl2b | ENSG00000083307 |
| ENSDARG00000045955 | prlrb | ENSG00000113494 |
| ENSDARG00000077532 | pign | ENSG00000197563 |
| ENSDARG00000098330 | spef2 | ENSG00000152582 |
| ENSDARG00000052408 | mgat2 | ENSG00000168282 |
| ENSDARG00000013667 | tbck | ENSG00000145348 |
| ENSDARG00000053502 | cryaa | ENSG00000160202 |
| ENSDARG00000079267 | FERMT3 (2 of 2) | ENSG00000149781 |
| ENSDARG00000071095 | abi3bpb | ENSG00000154175 |
| ENSDARG00000096932 | si:ch211-37e10.2 | ENSG00000165973 |
| ENSDARG00000100446 | si:ch211-286o17. | ENSG00000174059 |
| ENSDARG00000071594 | TIMM21 | ENSG00000075336 |
| ENSDARG00000016749 | slco1c1 | ENSG00000139155 |
| ENSDARG00000069009 | si:ch211-147a11.: | ENSG00000198912 |
| ENSDARG00000058656 | desma | ENSG00000175084 |
| ENSDARG00000070394 | uroc1 | ENSG00000159650 |
| ENSDARG00000057089 | ftsj1 | ENSG00000068438 |
| ENSDARG00000058178 | zdhhc20b | ENSG00000180776 |
| ENSDARG00000037064 | galca | ENSG00000054983 |
| ENSDARG00000098761 | rgs2 | ENSG00000116741 |
| ENSDARG00000092976 | si:ch211-127i16.2 | ENSG00000069535 |
| ENSDARG00000070229 | C11H1orf116 (1 c | ENSG00000182795 |
| ENSDARG00000090108 | DBNDD1 | ENSG00000003249 |
| ENSDARG00000017773 | slc16a12a | ENSG00000152779 |
| ENSDARG00000098989 | PLA2G3 (2 of 2) | ENSG00000100078 |
| ENSDARG00000054968 | cd40 | ENSG00000028137 |
| ENSDARG00000077547 | tmlhe | ENSG00000185973 |
| ENSDARG00000055632 | smtnl | ENSG00000188176 |
| ENSDARG00000069792 | crygm2d5 | ENSG00000118231 |
| ENSDARG00000078829 | ccdc149a | ENSG00000181982 |
| ENSDARG00000099524 | gbe1b | ENSG00000114480 |
| ENSDARG00000006514 | her6 | ENSG00000114315 |
| ENSDARG00000104370 | esm1 | ENSG00000164283 |
| ENSDARG00000029168 | ppfibp2b | ENSG00000166387 |
| ENSDARG00000029290 | stx11b.1 | ENSG00000135604 |

| | | |
|---|---|---|
| ENSDARG00000091996 | si:ch211-117m20. | ENSG00000134812 |
| ENSDARG00000055589 | s100t | ENSG00000143536 |
| ENSDARG00000055278 | cfb | ENSG00000243649 |
| ENSDARG00000056407 | irf8 | ENSG00000140968 |
| ENSDARG00000101878 | kdf1b | ENSG00000175707 |
| ENSDARG00000075942 | FRMD5 (1 of 2) | ENSG00000171877 |
| ENSDARG00000086879 | LINS1 | ENSG00000140471 |
| ENSDARG00000042725 | cebpb | ENSG00000172216 |
| ENSDARG00000089043 | ptpn6 | ENSG00000111679 |
| ENSDARG00000086881 | IER2 (1 of 2) | ENSG00000160888 |
| ENSDARG00000075215 | vetz | ENSG00000028203 |
| ENSDARG00000019496 | esd | ENSG00000139684 |
| ENSDARG00000078130 | zgc:193593 | ENSG00000104714 |
| ENSDARG00000038300 | rnf34a | ENSG00000170633 |
| ENSDARG00000026444 | apoob | ENSG00000184831 |
| ENSDARG00000017034 | sqrdl | ENSG00000137767 |
| ENSDARG00000055974 | tpmt.1 | ENSG00000137364 |
| ENSDARG00000032618 | mgst1.1 | ENSG00000008394 |
| ENSDARG00000036704 | tmem18 | ENSG00000151353 |
| ENSDARG00000054641 | fam46ab | ENSG00000112773 |
| ENSDARG00000015790 | atp1b3a | ENSG00000069849 |
| ENSDARG00000037358 | polr3e | ENSG00000058600 |
| ENSDARG00000075337 | lars2 | ENSG00000011376 |
| ENSDARG00000038941 | ppp1r1c | ENSG00000150722 |
| ENSDARG00000034941 | si:dkey-224e22.2 | ENSG00000149328 |
| ENSDARG00000045257 | decr2 | ENSG00000242612 |
| ENSDARG00000001313 | g2e3 | ENSG00000092140 |
| ENSDARG00000062959 | | ENSG00000173926 |
| ENSDARG00000101660 | sntg2 | ENSG00000172554 |
| ENSDARG00000060316 | cish | ENSG00000114737 |
| ENSDARG00000102185 | si:ch211-199f5.1 | ENSG00000136099 |
| ENSDARG00000104531 | zcchc4 | ENSG00000168228 |
| ENSDARG00000098249 | asmt | ENSG00000196433 |
| ENSDARG00000062633 | cadm2b | ENSG00000175161 |
| ENSDARG00000074248 | tmem237b | ENSG00000155755 |
| ENSDARG00000057105 | nle1 | ENSG00000073536 |
| ENSDARG00000095322 | FKBPL | ENSG00000204315 |
| ENSDARG00000044751 | ddt | ENSG00000099974 |
| ENSDARG00000039309 | hook2 | ENSG00000095066 |
| ENSDARG00000037895 | ramp2 | ENSG00000131477 |
| ENSDARG00000032116 | cnot7 | ENSG00000198791 |
| ENSDARG00000043093 | mpeg1.2 | ENSG00000197629 |
| ENSDARG00000069737 | pou4f2 | ENSG00000151615 |
| ENSDARG00000087373 | dnah5 | ENSG00000039139 |
| ENSDARG00000019861 | fgl2a | ENSG00000127951 |
| ENSDARG00000043562 | zgc:65997 | ENSG00000100867 |
| ENSDARG00000012039 | ttc26 | ENSG00000105948 |
| ENSDARG00000004037 | cog2 | ENSG00000135775 |
| ENSDARG00000077544 | toe1 | ENSG00000132773 |
| ENSDARG00000091754 | CHCHD5 | ENSG00000125611 |
| ENSDARG00000017985 | C1H4orf33 | ENSG00000151470 |
| ENSDARG00000101126 | atad5b | ENSG00000176208 |
| ENSDARG00000074226 | zgc:158398 | ENSG00000149932 |
| ENSDARG00000096932 | si:ch211-37e10.2 | ENSG00000165973 |
| ENSDARG00000043287 | atg101 | ENSG00000123395 |
| ENSDARG00000053559 | tspan3b | ENSG00000140391 |
| ENSDARG00000077804 | hgsnat | ENSG00000165102 |
| ENSDARG00000018002 | eci1 | ENSG00000167969 |
| ENSDARG00000098295 | efna3b | ENSG00000143590 |
| ENSDARG00000035521 | sfrp1a | ENSG00000104332 |
| ENSDARG00000008105 | mettl2a | ENSG00000087995 |
| ENSDARG00000054606 | mrpl1 | ENSG00000169288 |
| ENSDARG00000010183 | VWA5A (1 of 4) | ENSG00000055957 |
| ENSDARG00000059885 | frmd3 | ENSG00000172159 |
| ENSDARG00000042489 | prkrira | ENSG00000137492 |
| ENSDARG00000004218 | rnd1l | ENSG00000172602 |
| ENSDARG00000043719 | c3a.6 | ENSG00000125730 |
| ENSDARG00000040214 | scrt1b | ENSG00000261678 |
| ENSDARG00000078775 | RASA2 | ENSG00000155903 |
| ENSDARG00000060383 | mmaa | ENSG00000151611 |
| ENSDARG00000102302 | WDR48 (2 of 3) | ENSG00000114742 |
| ENSDARG00000013524 | cyp2ae1 | ENSG00000138115 |
| ENSDARG00000044318 | itgb7 | ENSG00000139626 |
| ENSDARG00000056642 | hdac9b | ENSG00000048052 |
| ENSDARG00000069499 | spag1b | ENSG00000104450 |
| ENSDARG00000040705 | glsb | ENSG00000115419 |
| ENSDARG00000079483 | CLIP1 (2 of 2) | ENSG00000130779 |
| ENSDARG00000052537 | cptp | ENSG00000182327 |
| ENSDARG00000013860 | ptpmt1 | ENSG00000110536 |
| ENSDARG00000036245 | WIPF3 | ENSG00000122574 |
| ENSDARG00000052624 | spryd7a | ENSG00000123178 |
| ENSDARG00000016139 | mkks | ENSG00000125863 |
| ENSDARG00000042045 | hmg20b | ENSG00000064961 |
| ENSDARG00000037954 | tnnt1 | ENSG00000105048 |
| ENSDARG00000032571 | il20ra | ENSG00000016402 |
| ENSDARG00000080021 | zmat3 | ENSG00000172667 |
| ENSDARG00000051956 | isca1 | ENSG00000135070 |
| ENSDARG00000053097 | hsf2 | ENSG00000025156 |
| ENSDARG00000062103 | ankrd24 | ENSG00000089847 |
| ENSDARG00000059202 | tspan2b | ENSG00000134198 |
| ENSDARG00000059438 | galnt18b | ENSG00000110328 |
| ENSDARG00000078272 | CABP7 (2 of 2) | ENSG00000100314 |
| ENSDARG00000029936 | grtp1a | ENSG00000139835 |
| ENSDARG00000024479 | pigk | ENSG00000142892 |
| ENSDARG00000026753 | grik4 | ENSG00000149403 |
| ENSDARG00000052625 | fkbp1b | ENSG00000119782 |
| ENSDARG00000044304 | prrc1 | ENSG00000164244 |
| ENSDARG00000055754 | smc1a | ENSG00000072501 |
| ENSDARG00000019596 | nrd1a | ENSG00000078618 |
| ENSDARG00000036061 | ogfod1 | ENSG00000087263 |
| ENSDARG00000104936 | si:ch211-225h24.: | ENSG00000180190 |

| | | |
|---|---|---|
| ENSDARG00000099390 | lama2 | ENSG00000196569 |
| ENSDARG00000006181 | hif1aa | ENSG00000100644 |
| ENSDARG00000051735 | si:ch211-113a14.: | ENSG00000187166 |
| ENSDARG00000022918 | tbcc | ENSG00000124659 |
| ENSDARG00000094677 | MPDZ | ENSG00000107186 |
| ENSDARG00000087040 | rhbdl1 | ENSG00000103269 |
| ENSDARG00000078829 | ccdc149a | ENSG00000181982 |
| ENSDARG00000045555 | sco2 | ENSG00000284194 |
| ENSDARG00000102097 | nfkbiz | ENSG00000144802 |
| ENSDARG00000068280 | GRB14 | ENSG00000115290 |
| ENSDARG00000074842 | snx8a | ENSG00000106266 |
| ENSDARG00000076017 | tmem69 | ENSG00000159596 |
| ENSDARG00000071567 | TSTD1 | ENSG00000215845 |
| ENSDARG00000007561 | cdh23 | ENSG00000107736 |
| ENSDARG00000058822 | wnt3a | ENSG00000154342 |
| ENSDARG00000078882 | SLC22A31 | ENSG00000259803 |
| ENSDARG00000104453 | flt4 | ENSG00000037280 |
| ENSDARG00000068344 | mmab | ENSG00000139428 |
| ENSDARG00000037281 | fgg | ENSG00000171557 |
| ENSDARG00000071065 | g6pd | ENSG00000160211 |
| ENSDARG00000104501 | si:dkey-108k21.9 | ENSG00000184678 |
| ENSDARG00000061737 | ece1 | ENSG00000117298 |
| ENSDARG00000088844 | deaf1 | ENSG00000177030 |
| ENSDARG00000070148 | cygb2 | ENSG00000161544 |
| ENSDARG00000035005 | bag2 | ENSG00000112208 |
| ENSDARG00000088181 | mpped1 | ENSG00000186732 |
| ENSDARG00000036139 | lctla | ENSG00000188501 |
| ENSDARG00000020239 | lpin1 | ENSG00000134324 |
| ENSDARG00000043431 | b3gntl1 | ENSG00000175711 |
| ENSDARG00000075673 | arhgap21b | ENSG00000107863 |
| ENSDARG00000043858 | cdk19 | ENSG00000155111 |
| ENSDARG00000042816 | mmp9 | ENSG00000100985 |
| ENSDARG00000039931 | slc25a33 | ENSG00000171612 |
| ENSDARG00000045159 | zdhhc4 | ENSG00000136247 |
| ENSDARG00000077553 | CABZ01067868.1 | ENSG00000125249 |
| ENSDARG00000035161 | pkdcca | ENSG00000162878 |
| ENSDARG00000099534 | CABZ01066926.1 | ENSG00000012963 |
| ENSDARG00000040280 | FAM84B (2 of 2) | ENSG00000168672 |
| ENSDARG00000062949 | nt5dc3 | ENSG00000111696 |
| ENSDARG00000100115 | dnah1 | ENSG00000114841 |
| ENSDARG00000088764 | lyplal1 | ENSG00000143353 |
| ENSDARG00000031845 | mboat2a | ENSG00000143797 |
| ENSDARG00000038901 | tmem256 | ENSG00000205544 |
| ENSDARG00000030753 | prrg4 | ENSG00000135378 |
| ENSDARG00000058522 | shq1 | ENSG00000144736 |
| ENSDARG00000014333 | ormdl1 | ENSG00000128699 |
| ENSDARG00000013842 | tbc1d19 | ENSG00000109680 |
| ENSDARG00000028396 | fkbp5 | ENSG00000096060 |
| ENSDARG00000044627 | alg8 | ENSG00000159063 |
| ENSDARG00000076862 | fam198a | ENSG00000144649 |
| ENSDARG00000053380 | fam189a1 | ENSG00000104059 |
| ENSDARG00000056648 | ext2 | ENSG00000151348 |
| ENSDARG00000070119 | ctc1 | ENSG00000178971 |
| ENSDARG00000077870 | taf9 | ENSG00000273841 |
| ENSDARG00000099200 | zgc:123103 | ENSG00000135047 |
| ENSDARG00000056642 | hdac9b | ENSG00000048052 |
| ENSDARG00000070485 | apitd1 | ENSG00000241563 |
| ENSDARG00000101800 | PSTK | ENSG00000179988 |
| ENSDARG00000098401 | fam136a | ENSG00000035141 |
| ENSDARG00000063161 | ppwd1 | ENSG00000113593 |
| ENSDARG00000036291 | nucb2b | ENSG00000070081 |
| ENSDARG00000098461 | si:dkey-11f4.14 | ENSG00000117091 |
| ENSDARG00000060124 | timm17b | ENSG00000126768 |
| ENSDARG00000074617 | wfs1b | ENSG00000109501 |
| ENSDARG00000102846 | trmt13 | ENSG00000122435 |
| ENSDARG00000021480 | olfml2bb | ENSG00000162745 |
| ENSDARG00000020595 | gnl3l | ENSG00000130119 |
| ENSDARG00000075687 | anapc1 | ENSG00000153107 |
| ENSDARG00000105077 | cecr5 | ENSG00000069998 |
| ENSDARG00000004171 | rabac1 | ENSG00000105404 |
| ENSDARG00000104370 | esm1 | ENSG00000164283 |
| ENSDARG00000089976 | spcs3 | ENSG00000129128 |
| ENSDARG00000039482 | calhm2 | ENSG00000138172 |
| ENSDARG00000057437 | apodb | ENSG00000189058 |
| ENSDARG00000031098 | timm50 | ENSG00000105197 |
| ENSDARG00000075923 | TXNRD2 (1 of 2) | ENSG00000184470 |
| ENSDARG00000099708 | cacna1hb | ENSG00000196557 |
| ENSDARG00000023316 | sim1a | ENSG00000112246 |
| ENSDARG00000075948 | slco4a1 | ENSG00000101187 |
| ENSDARG00000063574 | mtg2 | ENSG00000101181 |
| ENSDARG00000075016 | apobb.2 | ENSG00000084674 |
| ENSDARG00000079808 | si:ch73-208g10.1 | ENSG00000076662 |
| ENSDARG00000057644 | adam8b | ENSG00000151651 |
| ENSDARG00000062651 | zc3hc1 | ENSG00000091732 |
| ENSDARG00000023820 | faxdc2 | ENSG00000170271 |
| ENSDARG00000100378 | TM4SF19 | ENSG00000145107 |
| ENSDARG00000071203 | sptssa | ENSG00000165389 |
| ENSDARG00000023672 | timm8a | ENSG00000126953 |
| ENSDARG00000027353 | zmym2 | ENSG00000121741 |
| ENSDARG00000060196 | lhpp | ENSG00000107902 |
| ENSDARG00000075393 | spock2 | ENSG00000107742 |
| ENSDARG00000099801 | topaz1 | ENSG00000173769 |
| ENSDARG00000004158 | tmem63c | ENSG00000165548 |
| ENSDARG00000042025 | C9H2orf49 | ENSG00000135974 |
| ENSDARG00000059856 | alkbh2 | ENSG00000189046 |
| ENSDARG00000053318 | lrrc6 | ENSG00000129295 |
| ENSDARG00000077920 | CSMD3 | ENSG00000164796 |
| ENSDARG00000103114 | ndufa4l2a | ENSG00000185633 |
| ENSDARG00000045304 | mtfr1 | ENSG00000066855 |
| ENSDARG00000046133 | b3galnt2 | ENSG00000162885 |
| ENSDARG00000041910 | znf512 | ENSG00000243943 |

| | | |
|---|---|---|
| ENSDARG00000036291 | nucb2b | ENSG00000070081 |
| ENSDARG00000043443 | kcnj13 | ENSG00000115474 |
| ENSDARG00000052176 | C3H19orf66 | ENSG00000130813 |
| ENSDARG00000029307 | C24H18orf8 | ENSG00000141452 |
| ENSDARG00000079249 | OGDHL | ENSG00000197444 |
| ENSDARG00000055825 | celsr3 | ENSG00000008300 |
| ENSDARG00000010407 | ppp4r4 | ENSG00000119698 |
| ENSDARG00000100280 | znfx1 | ENSG00000124201 |
| ENSDARG00000099315 | nsdhl | ENSG00000147383 |
| ENSDARG00000060741 | SYT17 | ENSG00000103528 |
| ENSDARG00000030630 | mfsd2aa | ENSG00000168389 |
| ENSDARG00000099385 | LSP1 (2 of 2) | ENSG00000130592 |
| ENSDARG00000044437 | cdca5 | ENSG00000146670 |
| ENSDARG00000022845 | lias | ENSG00000121897 |
| ENSDARG00000076179 | ngef | ENSG00000066248 |
| ENSDARG00000074633 | gpr35.1 | ENSG00000178623 |
| ENSDARG00000056379 | si:ch211-154o6.6 | ENSG00000152672 |
| ENSDARG00000045983 | zbtb43 | ENSG00000169155 |
| ENSDARG00000041589 | adprhl1 | ENSG00000153531 |
| ENSDARG00000016483 | baiap2l2a | ENSG00000128298 |
| ENSDARG00000095157 | TMEM14A | ENSG00000096092 |
| ENSDARG00000031202 | alg11 | ENSG00000253710 |
| ENSDARG00000058287 | gpalpp1 | ENSG00000133114 |
| ENSDARG00000059634 | thumpd3 | ENSG00000134077 |
| ENSDARG00000029457 | cacna1sa | ENSG00000081248 |
| ENSDARG00000018688 | elk3 | ENSG00000111145 |
| ENSDARG00000037259 | ndufb6 | ENSG00000165264 |
| ENSDARG00000087916 | tmem240a | ENSG00000205090 |
| ENSDARG00000013251 | hsf4 | ENSG00000102878 |
| ENSDARG00000076508 | grm8b | ENSG00000179603 |
| ENSDARG00000094605 | si:ch211-154e10.1 | ENSG00000138835 |
| ENSDARG00000052769 | glrbb | ENSG00000109738 |
| ENSDARG00000102375 | BDP1 (2 of 2) | ENSG00000145734 |
| ENSDARG00000034174 | mrrf | ENSG00000148187 |
| ENSDARG00000020976 | ubald1b | ENSG00000153443 |
| ENSDARG00000097613 | igfn1.3 | ENSG00000179057 |
| ENSDARG00000062084 | clcn1a | ENSG00000188037 |
| ENSDARG00000003167 | mlst8 | ENSG00000167965 |
| ENSDARG00000059263 | hoxd12a | ENSG00000170178 |
| ENSDARG00000087489 | FAM124B | ENSG00000124019 |
| ENSDARG00000053853 | slc13a2 | ENSG00000007216 |
| ENSDARG00000004794 | ccdc146 | ENSG00000135205 |
| ENSDARG00000016457 | irf9 | ENSG00000213928 |
| ENSDARG00000070463 | e2f3 | ENSG00000112242 |
| ENSDARG00000006577 | nadka | ENSG00000008130 |
| ENSDARG00000003869 | decr1 | ENSG00000104325 |
| ENSDARG00000102050 | CABZ01069335.1 | ENSG00000055732 |
| ENSDARG00000075380 | kri1 | ENSG00000129347 |
| ENSDARG00000021924 | hsp70.3 | ENSG00000126803 |
| ENSDARG00000101322 | tfr1a | ENSG00000072274 |
| ENSDARG00000073870 | gdpd2 | ENSG00000130055 |
| ENSDARG00000000151 | thraa | ENSG00000126351 |
| ENSDARG00000003022 | limk1b | ENSG00000106683 |
| ENSDARG00000071573 | lsm5 | ENSG00000106355 |
| ENSDARG00000062552 | lpar6a | ENSG00000139679 |
| ENSDARG00000005716 | olfml2ba | ENSG00000162745 |
| ENSDARG00000044405 | ttc4 | ENSG00000243725 |
| ENSDARG00000028053 | bmp1b | ENSG00000168487 |
| ENSDARG00000042659 | thyn1 | ENSG00000151500 |
| ENSDARG00000089043 | ptpn6 | ENSG00000111679 |
| ENSDARG00000041011 | gemin7 | ENSG00000142252 |
| ENSDARG00000041736 | hsdl1 | ENSG00000103160 |
| ENSDARG00000061977 | ppfibp2a | ENSG00000166387 |
| ENSDARG00000021208 | serpind1 | ENSG00000099937 |
| ENSDARG00000101190 | cyp2k8 | ENSG00000108242 |
| ENSDARG00000016649 | akr7a3 | ENSG00000211454 |
| ENSDARG00000071877 | dhrs7cb | ENSG00000184544 |
| ENSDARG00000068934 | cyp1b1 | ENSG00000138061 |
| ENSDARG00000012981 | osbpl7 | ENSG00000006025 |
| ENSDARG00000088347 | sp1 | ENSG00000185591 |
| ENSDARG00000059558 | tecta | ENSG00000109927 |
| ENSDARG00000059116 | heatr5b | ENSG00000008869 |
| ENSDARG00000100352 | aglb | ENSG00000162688 |
| ENSDARG00000028173 | slc4a2a | ENSG00000164889 |
| ENSDARG00000040466 | vil1 | ENSG00000127831 |
| ENSDARG00000012829 | asah2 | ENSG00000188611 |
| ENSDARG00000001712 | si:ch211-247n2.1 | ENSG00000145936 |
| ENSDARG00000078076 | nab2 | ENSG00000166886 |
| ENSDARG00000021509 | tab2 | ENSG00000055208 |
| ENSDARG00000102185 | si:ch211-199f5.1 | ENSG00000136099 |
| ENSDARG00000019265 | ZBTB39 | ENSG00000166860 |
| ENSDARG00000006865 | glra4a | ENSG00000188828 |
| ENSDARG00000056438 | her9 | ENSG00000188290 |
| ENSDARG00000038964 | traf4b | ENSG00000076604 |
| ENSDARG00000026736 | il11ra | ENSG00000096996 |
| ENSDARG00000099348 | ITGA6 (4 of 4) | ENSG00000091409 |
| ENSDARG00000062579 | kremen1 | ENSG00000183762 |
| ENSDARG00000099178 | ABCD2 | ENSG00000173208 |
| ENSDARG00000070917 | kitlga | ENSG00000049130 |
| ENSDARG00000078326 | arhgap10 | ENSG00000071205 |
| ENSDARG00000075974 | si:dkey-258f14.3 | ENSG00000130164 |
| ENSDARG00000063435 | trpm4b.2 | ENSG00000130529 |
| ENSDARG00000002764 | miox | ENSG00000100253 |
| ENSDARG00000032340 | serhl | ENSG00000183569 |
| ENSDARG00000042722 | blnk | ENSG00000095585 |
| ENSDARG00000063634 | hpse | ENSG00000173083 |
| ENSDARG00000074253 | olig3 | ENSG00000177468 |
| ENSDARG00000008414 | exoc3l2a | ENSG00000283632 |
| ENSDARG00000057273 | alox5a | ENSG00000012779 |
| ENSDARG00000103476 | trpm1b | ENSG00000134160 |
| ENSDARG00000056938 | kera | ENSG00000139330 |
| ENSDARG00000060002 | ogg1 | ENSG00000114026 |
| ENSDARG00000057562 | polr3c | ENSG00000186141 |
| ENSDARG00000005619 | nek2 | ENSG00000117650 |
| ENSDARG00000095034 | C24H8orf34 | ENSG00000165084 |
| ENSDARG00000070399 | alg2 | ENSG00000119523 |
| ENSDARG00000037000 | fkbp11 | ENSG00000134285 |
| ENSDARG00000075016 | apobb.2 | ENSG00000084674 |
| ENSDARG00000016483 | baiap2l2a | ENSG00000128298 |
| ENSDARG00000030108 | sec22ba | ENSG00000265808 |
| ENSDARG00000086735 | SAMD4B | ENSG00000179134 |
| ENSDARG00000026902 | cryzl1 | ENSG00000205758 |
| ENSDARG00000086691 | dnajc12 | ENSG00000108176 |
| ENSDARG00000055223 | ablim2 | ENSG00000163995 |
| ENSDARG00000098600 | pithd1 | ENSG00000057757 |
| ENSDARG00000077863 | mettl22 | ENSG00000067365 |
| ENSDARG00000057021 | vps36 | ENSG00000136100 |
| ENSDARG00000079414 | sez6b | ENSG00000063015 |
| ENSDARG00000039265 | arhgap4a | ENSG00000089820 |
| ENSDARG00000095614 | cacna1eb | ENSG00000198216 |
| ENSDARG00000021555 | park2 | ENSG00000185345 |
| ENSDARG00000058327 | neu3.3 | ENSG00000162139 |
| ENSDARG00000030761 | syt16 | ENSG00000139973 |
| ENSDARG00000089172 | | ENSG00000173482 |
| ENSDARG00000024381 | msto1 | ENSG00000125459 |
| ENSDARG00000038964 | traf4b | ENSG00000076604 |
| ENSDARG00000057984 | eps8l1 | ENSG00000131037 |
| ENSDARG00000054821 | nubpl | ENSG00000151413 |
| ENSDARG00000034855 | ccnb3 | ENSG00000147082 |
| ENSDARG00000042557 | rnf144ab | ENSG00000151692 |
| ENSDARG00000057606 | arl4ca | ENSG00000188042 |
| ENSDARG00000070085 | mettl6 | ENSG00000206562 |
| ENSDARG00000086998 | NNMT | ENSG00000166741 |
| ENSDARG00000068939 | XAF1 | ENSG00000132530 |
| ENSDARG00000062363 | phex | ENSG00000102174 |
| ENSDARG00000010029 | spon1a | ENSG00000262655 |
| ENSDARG00000005559 | pou4f1 | ENSG00000152192 |
| ENSDARG00000056400 | helt | ENSG00000187821 |
| ENSDARG00000003077 | si:ch73-111m19.2 | ENSG00000233276 |
| ENSDARG00000036162 | hnrnpa0b | ENSG00000170144 |
| ENSDARG00000060875 | pde1a | ENSG00000115252 |
| ENSDARG00000055331 | aldh9a1a.2 | ENSG00000143149 |
| ENSDARG00000104274 | TMEM200B | ENSG00000253304 |
| ENSDARG00000015355 | fosl1a | ENSG00000175592 |
| ENSDARG00000090145 | tmem240b | ENSG00000205090 |
| ENSDARG00000067752 | pigl | ENSG00000108474 |
| ENSDARG00000090871 | si:dkey-210j14.4 | ENSG00000165512 |
| ENSDARG00000067509 | slc24a4b | ENSG00000140090 |
| ENSDARG00000044380 | rbmx2 | ENSG00000134597 |
| ENSDARG00000104217 | gpc2 | ENSG00000213420 |
| ENSDARG00000039325 | pmepa1 | ENSG00000124225 |
| ENSDARG00000044142 | acss1 | ENSG00000154930 |
| ENSDARG00000079065 | ANKRD50 (2 of 2) | ENSG00000151458 |
| ENSDARG00000071429 | tdo2a | ENSG00000151790 |
| ENSDARG00000009949 | FNDC4 (1 of 2) | ENSG00000115226 |
| ENSDARG00000013292 | tmem179 | ENSG00000258986 |
| ENSDARG00000040442 | prtgb | ENSG00000166450 |
| ENSDARG00000077664 | zgc:110239 | ENSG00000135047 |
| ENSDARG00000069383 | cnga4 | ENSG00000132259 |
| ENSDARG00000060554 | mut | ENSG00000146085 |
| ENSDARG00000059916 | hyal1 | ENSG00000114378 |
| ENSDARG00000104186 | fbxo43 | ENSG00000156509 |
| ENSDARG00000008840 | hmbsa | ENSG00000256269 |
| ENSDARG00000095022 | thsd7ba | ENSG00000144229 |
| ENSDARG00000051874 | stra6 | ENSG00000137868 |
| ENSDARG00000076296 | triqk | ENSG00000205133 |
| ENSDARG00000040116 | znf366 | ENSG00000178175 |
| ENSDARG00000103703 | CLDN23 (3 of 3) | ENSG00000253958 |
| ENSDARG00000011727 | b9d1 | ENSG00000108641 |
| ENSDARG00000024109 | naa40 | ENSG00000110583 |
| ENSDARG00000007244 | acp2 | ENSG00000134575 |
| ENSDARG00000021220 | hmgcl | ENSG00000117305 |
| ENSDARG00000016480 | slc17a7a | ENSG00000104888 |
| ENSDARG00000062884 | map3k2 | ENSG00000169967 |
| ENSDARG00000023820 | faxdc2 | ENSG00000170271 |
| ENSDARG00000098780 | nsmce1 | ENSG00000169189 |
| ENSDARG00000099133 | UNC5C (2 of 2) | ENSG00000182168 |
| ENSDARG00000018441 | heg | ENSG00000173706 |
| ENSDARG00000038789 | tmem53 | ENSG00000126106 |
| ENSDARG00000034423 | sncga | ENSG00000173267 |
| ENSDARG00000027070 | acbd4 | ENSG00000181513 |
| ENSDARG00000059816 | C1H19orf53 | ENSG00000104979 |
| ENSDARG00000023683 | cacna1fb | ENSG00000102001 |
| ENSDARG00000101341 | C11H19orf24 | ENSG00000228300 |
| ENSDARG00000077202 | mocs2 | ENSG00000164172 |
| ENSDARG00000090195 | mtx3 | ENSG00000177034 |
| ENSDARG00000061348 | adat1 | ENSG00000065457 |
| ENSDARG00000036930 | tbx18 | ENSG00000112837 |
| ENSDARG00000045790 | C4H12orf66 | ENSG00000174206 |
| ENSDARG00000039577 | ptk2bb | ENSG00000120899 |
| ENSDARG00000042159 | RFESD (1 of 2) | ENSG00000175449 |
| ENSDARG00000069427 | nkain2 | ENSG00000188580 |
| ENSDARG00000100559 | tor2a | ENSG00000160404 |
| ENSDARG00000102937 | hspbp1 | ENSG00000133265 |
| ENSDARG00000020730 | smpd4 | ENSG00000136699 |
| ENSDARG00000046081 | bloc1s2 | ENSG00000196072 |
| ENSDARG00000060871 | mctp1b | ENSG00000175471 |
| ENSDARG00000012981 | osbpl7 | ENSG00000006025 |
| ENSDARG00000098209 | SIMC1 | ENSG00000170085 |
| ENSDARG00000038006 | osr2 | ENSG00000164920 |
| ENSDARG00000045153 | slc16a8 | ENSG00000100156 |
| ENSDARG00000104179 | rab11fip5a | ENSG00000135631 |
| ENSDARG00000057890 | TTC31 | ENSG00000115282 |
| ENSDARG00000031248 | enkur | ENSG00000151023 |
| ENSDARG00000023989 | pef1 | ENSG00000162517 |
| ENSDARG00000088463 | FO904873.1 | ENSG00000162129 |
| ENSDARG00000052617 | raver1 | ENSG00000161847 |
| ENSDARG00000002483 | WNT7A (1 of 2) | ENSG00000154764 |
| ENSDARG00000038567 | tsr3 | ENSG00000007520 |
| ENSDARG00000031796 | ache | ENSG00000087085 |
| ENSDARG00000005374 | tubgcp4 | ENSG00000137822 |
| ENSDARG00000054610 | coro1a | ENSG00000102879 |
| ENSDARG00000008127 | pcdh15b | ENSG00000150275 |
| ENSDARG00000099732 | gfra1a | ENSG00000151892 |
| ENSDARG00000101628 | ascl1b | ENSG00000183734 |
| ENSDARG00000045175 | pofut2 | ENSG00000186866 |
| ENSDARG00000010981 | zdhhc3a | ENSG00000163812 |
| ENSDARG00000070081 | rspo3 | ENSG00000146374 |
| ENSDARG00000039020 | fbxo5 | ENSG00000112029 |
| ENSDARG00000057830 | dnmt3bb.2 | ENSG00000142182 |
| ENSDARG00000079815 | lyrm7 | ENSG00000186687 |
| ENSDARG00000043012 | napgb | ENSG00000134265 |
| ENSDARG00000103464 | pggt1b | ENSG00000164219 |
| ENSDARG00000033832 | ampd1 | ENSG00000116748 |
| ENSDARG00000035406 | ttc33 | ENSG00000113638 |
| ENSDARG00000058587 | ccdc79 | ENSG00000249961 |
| ENSDARG00000062103 | ankrd24 | ENSG00000089847 |
| ENSDARG00000032469 | ampd3b | ENSG00000133805 |
| ENSDARG00000040707 | CHCHD4 (2 of 2) | ENSG00000163528 |
| ENSDARG00000062171 | olfml3b | ENSG00000116774 |
| ENSDARG00000032816 | tm7sf2 | ENSG00000149809 |
| ENSDARG00000060394 | mtfr2 | ENSG00000146410 |
| ENSDARG00000031890 | tcp11l1 | ENSG00000176148 |
| ENSDARG00000067593 | SCO1 | ENSG00000133028 |
| ENSDARG00000017320 | f11r.1 | ENSG00000158769 |
| ENSDARG00000090408 | ERBB4 (3 of 3) | ENSG00000178568 |
| ENSDARG00000025034 | syngr3b | ENSG00000127561 |
| ENSDARG00000034174 | mrrf | ENSG00000148187 |
| ENSDARG00000028367 | sult2st3 | ENSG00000088002 |
| ENSDARG00000099412 | bcan | ENSG00000132692 |
| ENSDARG00000056609 | atad1b | ENSG00000138138 |
| ENSDARG00000034895 | tgfb1b | ENSG00000105329 |
| ENSDARG00000054950 | eef2kmt | ENSG00000118894 |
| ENSDARG00000036894 | AIMP1 (1 of 2) | ENSG00000164022 |
| ENSDARG00000052247 | alkbh4 | ENSG00000160993 |
| ENSDARG00000075639 | si:ch211-89o9.6 | ENSG00000240747 |
| ENSDARG00000041699 | piwil1 | ENSG00000125207 |
| ENSDARG00000090230 | tmem108 | ENSG00000144868 |
| ENSDARG00000060944 | rnf217 | ENSG00000146373 |
| ENSDARG00000068698 | psenen | ENSG00000188223 |
| ENSDARG00000009557 | wrap73 | ENSG00000116213 |
| ENSDARG00000067672 | card9 | ENSG00000187796 |
| ENSDARG00000077817 | cxxc4 | ENSG00000168772 |
| ENSDARG00000063646 | lnpa | ENSG00000144320 |
| ENSDARG00000071465 | lrrc39 | ENSG00000122477 |
| ENSDARG00000015807 | rab22a | ENSG00000124209 |
| ENSDARG00000002593 | slc45a2 | ENSG00000164175 |
| ENSDARG00000006526 | fn1b | ENSG00000115414 |
| ENSDARG00000041493 | ipo4 | ENSG00000196497 |
| ENSDARG00000021110 | poc1b | ENSG00000119599 |
| ENSDARG00000104244 | twsg1a | ENSG00000128791 |
| ENSDARG00000045598 | si:dkey-180p18.9 | ENSG00000085662 |
| ENSDARG00000101190 | cyp2k8 | ENSG00000108242 |
| ENSDARG00000017612 | pnpo | ENSG00000108439 |
| ENSDARG00000077465 | SORCS2 | ENSG00000184985 |
| ENSDARG00000020015 | mrps33 | ENSG00000090263 |
| ENSDARG00000039424 | snrpb2 | ENSG00000125870 |
| ENSDARG00000078853 | arhgef19 | ENSG00000142632 |
| ENSDARG00000028581 | jkamp | ENSG00000050130 |
| ENSDARG00000041864 | capn3a | ENSG00000092529 |
| ENSDARG00000045289 | gpr137c | ENSG00000180998 |
| ENSDARG00000069313 | oxa1l | ENSG00000155463 |
| ENSDARG00000024365 | crlf1a | ENSG00000006016 |
| ENSDARG00000097530 | XKR5 (1 of 2) | ENSG00000275591 |
| ENSDARG00000061354 | foxred2 | ENSG00000100350 |
| ENSDARG00000053344 | adgrd1 | ENSG00000111452 |
| ENSDARG00000002311 | fabp11b | ENSG00000147588 |
| ENSDARG00000020241 | icmt | ENSG00000116237 |
| ENSDARG00000100805 | FMN1 (3 of 4) | ENSG00000248905 |
| ENSDARG00000069662 | fgf11a | ENSG00000161958 |
| ENSDARG00000014057 | gabrr2b | ENSG00000111886 |
| ENSDARG00000083400 | SNORA73 | ENSG00000222145 |
| ENSDARG00000078650 | KCNA4 | ENSG00000182255 |
| ENSDARG00000052734 | hmgcra | ENSG00000113161 |
| ENSDARG00000058365 | hspb8 | ENSG00000152137 |
| ENSDARG00000020741 | fga | ENSG00000171560 |
| ENSDARG00000039980 | ppt1 | ENSG00000131238 |
| ENSDARG00000063001 | atp8a1 | ENSG00000124406 |
| ENSDARG00000021120 | ndc1 | ENSG00000058804 |
| ENSDARG00000070954 | hmx2 | ENSG00000188816 |
| ENSDARG00000071414 | C12H17orf62 | ENSG00000178927 |
| ENSDARG00000035028 | lpcat4 | ENSG00000176454 |
| ENSDARG00000007990 | wt1b | ENSG00000184937 |
| ENSDARG00000086550 | dbf4b | ENSG00000161692 |
| ENSDARG00000079571 | zgc:162161 | ENSG00000173214 |
| ENSDARG00000014373 | ddx4 | ENSG00000152670 |
| ENSDARG00000069501 | abhd11 | ENSG00000106077 |
| ENSDARG00000062579 | kremen1 | ENSG00000183762 |
| ENSDARG00000026796 | grm1a | ENSG00000152822 |
| ENSDARG00000079434 | si:ch211-208h7.4 | ENSG00000121210 |
| ENSDARG00000100558 | slbp | ENSG00000163950 |
| ENSDARG00000012271 | tmem206 | ENSG00000065600 |

| | | | | | |
|---|---|---|---|---|---|
| ENSDARG00000041429 | vwa9 | ENSG00000138614 | ENSDARG00000070522 | cacna1i | ENSG00000100346 |
| | | | ENSDARG00000045874 | yeats4 | ENSG00000127337 |
| | | | ENSDARG00000035273 | tmtc2b | ENSG00000179104 |
| | | | ENSDARG00000015747 | aarsd1 | ENSG00000108825 |
| | | | ENSDARG00000061228 | pfdn4 | ENSG00000101132 |
| | | | ENSDARG00000015355 | fosl1a | ENSG00000175592 |
| | | | ENSDARG00000005518 | slc5a9 | ENSG00000117834 |
| | | | ENSDARG00000063568 | syt7a | ENSG00000011347 |
| | | | ENSDARG00000102855 | neo1a | ENSG00000067141 |
| | | | ENSDARG00000023111 | plg | ENSG00000122194 |
| | | | ENSDARG00000086107 | MTERF1 | ENSG00000127989 |
| | | | ENSDARG00000102773 | cacna1da | ENSG00000157388 |
| | | | ENSDARG00000091061 | slc38a3b | ENSG00000188338 |
| | | | ENSDARG00000043635 | cmc2 | ENSG00000103121 |
| | | | ENSDARG00000060871 | mctp1b | ENSG00000175471 |
| | | | ENSDARG00000033757 | ncaph2 | ENSG00000025770 |
| | | | ENSDARG00000078781 | si:ch211-161c3.6 | ENSG00000137309 |
| | | | ENSDARG00000069954 | kcnq5a | ENSG00000185760 |
| | | | ENSDARG00000101726 | CES3 (3 of 4) | ENSG00000172828 |
| | | | ENSDARG00000035538 | atp6v0a2a | ENSG00000185344 |
| | | | ENSDARG00000079324 | ANO9 (2 of 2) | ENSG00000185101 |
| | | | ENSDARG00000031598 | calb1 | ENSG00000104327 |
| | | | ENSDARG00000102312 | CABZ01063402.1 | ENSG00000185313 |
| | | | ENSDARG00000059883 | trpv1 | ENSG00000187688 |
| | | | ENSDARG00000016457 | irf9 | ENSG00000213928 |
| | | | ENSDARG00000075980 | tmem125b | ENSG00000179178 |
| | | | ENSDARG00000045754 | mettl25 | ENSG00000127720 |
| | | | ENSDARG00000058260 | zgc:174160 | ENSG00000163507 |
| | | | ENSDARG00000074335 | boc | ENSG00000144857 |
| | | | ENSDARG00000037409 | tmem144a | ENSG00000164124 |
| | | | ENSDARG00000011272 | uqcc2 | ENSG00000137288 |
| | | | ENSDARG00000045748 | stab2 | ENSG00000136011 |
| | | | ENSDARG00000038010 | rac2 | ENSG00000128340 |
| | | | ENSDARG00000104149 | rraga | ENSG00000083750 |
| | | | ENSDARG00000006862 | kifap3b | ENSG00000075945 |
| | | | ENSDARG00000020645 | slc7a3a | ENSG00000165349 |
| | | | ENSDARG00000074623 | tbc1d31 | ENSG00000156787 |
| | | | ENSDARG00000005368 | mcamb | ENSG00000076706 |
| | | | ENSDARG00000091092 | zcchc10 | ENSG00000155329 |
| | | | ENSDARG00000000767 | spi1b | ENSG00000066336 |
| | | | ENSDARG00000100657 | egfl7 | ENSG00000172889 |
| | | | ENSDARG00000055893 | nmnat1 | ENSG00000139194 |
| | | | ENSDARG00000068787 | slc6a17 | ENSG00000197106 |
| | | | ENSDARG00000014939 | KCNN2 | ENSG00000080709 |
| | | | ENSDARG00000012194 | scp2a | ENSG00000116171 |
| | | | ENSDARG00000079353 | si:ch211-165d12. | ENSG00000173762 |
| | | | ENSDARG00000017049 | adsl | ENSG00000239900 |
| | | | ENSDARG00000105035 | anapc2 | ENSG00000176248 |
| | | | ENSDARG00000009208 | prkcda | ENSG00000163932 |
| | | | ENSDARG00000061796 | rasgrp4 | ENSG00000171777 |
| | | | ENSDARG00000063522 | bbs4 | ENSG00000140463 |
| | | | ENSDARG00000036626 | sema6ba | ENSG00000167680 |
| | | | ENSDARG00000079102 | PRRG3 | ENSG00000130032 |
| | | | ENSDARG00000055278 | cfb | ENSG00000243649 |
| | | | ENSDARG00000062929 | FAM83G (1 of 2) | ENSG00000188522 |
| | | | ENSDARG00000013057 | hoxb5a | ENSG00000120075 |
| | | | ENSDARG00000076997 | stxbp4 | ENSG00000166263 |
| | | | ENSDARG00000041565 | tnfaip1 | ENSG00000109079 |
| | | | ENSDARG00000040440 | snrpd2 | ENSG00000125743 |
| | | | ENSDARG00000053127 | helb | ENSG00000127311 |
| | | | ENSDARG00000059234 | mrps27 | ENSG00000113048 |
| | | | ENSDARG00000017255 | tmed1b | ENSG00000099203 |
| | | | ENSDARG00000059054 | pdk2b | ENSG00000005882 |
| | | | ENSDARG00000033735 | ncf1 | ENSG00000158517 |
| | | | ENSDARG00000090943 | CABZ01033205.2 | ENSG00000168955 |
| | | | ENSDARG00000102587 | mios | ENSG00000164654 |
| | | | ENSDARG00000060620 | lig4 | ENSG00000174405 |
| | | | ENSDARG00000038025 | cbx7a | ENSG00000100307 |
| | | | ENSDARG00000043986 | setd6 | ENSG00000103037 |
| | | | ENSDARG00000058561 | fbxo30a | ENSG00000118496 |
| | | | ENSDARG00000007429 | ndor1 | ENSG00000188566 |
| | | | ENSDARG00000010437 | fam46c | ENSG00000183508 |
| | | | ENSDARG00000077180 | slc37a4b | ENSG00000112337 |
| | | | ENSDARG00000044142 | acss1 | ENSG00000154930 |
| | | | ENSDARG00000062370 | bcl2l13 | ENSG00000099968 |
| | | | ENSDARG00000093546 | ms4a17a.2 | ENSG00000110077 |
| | | | ENSDARG00000036864 | slc34a2b | ENSG00000157765 |
| | | | ENSDARG00000070675 | fam3a | ENSG00000071889 |
| | | | ENSDARG00000062748 | pkn1b | ENSG00000123143 |
| | | | ENSDARG00000075962 | vav3b | ENSG00000134215 |
| | | | ENSDARG00000060366 | slc12a9 | ENSG00000146828 |
| | | | ENSDARG00000027159 | fbxo28 | ENSG00000143756 |
| | | | ENSDARG00000077339 | ptrh2 | ENSG00000141378 |
| | | | ENSDARG00000076328 | osbpl11 | ENSG00000144909 |
| | | | ENSDARG00000078318 | tmod1 | ENSG00000136842 |
| | | | ENSDARG00000036305 | phf23b | ENSG00000040633 |
| | | | ENSDARG00000059835 | zfyve27 | ENSG00000155256 |
| | | | ENSDARG00000035562 | mpdu1a | ENSG00000129255 |
| | | | ENSDARG00000070824 | ndufb5 | ENSG00000136521 |
| | | | ENSDARG00000087120 | slc5a8 | ENSG00000256870 |
| | | | ENSDARG00000012671 | inhbaa | ENSG00000122641 |
| | | | ENSDARG00000052703 | cisd2 | ENSG00000145354 |
| | | | ENSDARG00000076804 | ttyh1 | ENSG00000167614 |
| | | | ENSDARG00000091869 | zgc:162958 | ENSG00000167394 |
| | | | ENSDARG00000001897 | man2b1 | ENSG00000104774 |
| | | | ENSDARG00000098249 | asmt | ENSG00000196433 |
| | | | ENSDARG00000060946 | pip5kl1 | ENSG00000167103 |
| | | | ENSDARG00000023546 | kcnn1b | ENSG00000105642 |
| | | | ENSDARG00000095536 | TRIM62 (2 of 2) | ENSG00000116525 |
| | | | ENSDARG00000089032 | ZNF384 (2 of 2) | ENSG00000126746 |
| | | | ENSDARG00000025420 | fzd5 | ENSG00000163251 |

| | | |
|---|---|---|
| ENSDARG00000004539 | ptgs2a | ENSG00000073756 |
| ENSDARG00000034714 | esyt1a | ENSG00000139641 |
| ENSDARG00000006758 | FAM234B | ENSG00000084444 |
| ENSDARG00000057113 | c6 | ENSG00000039537 |
| ENSDARG00000101527 | IL1R2 (2 of 2) | ENSG00000115602 |
| ENSDARG00000076020 | pappa2 | ENSG00000116183 |
| ENSDARG00000013963 | mipb | ENSG00000135517 |
| ENSDARG00000030805 | znhit2 | ENSG00000174276 |
| ENSDARG00000074204 | commd6 | ENSG00000188243 |
| ENSDARG00000026484 | rab15 | ENSG00000139998 |
| ENSDARG00000045355 | tceb1a | ENSG00000154582 |
| ENSDARG00000098726 | rptor | ENSG00000141564 |
| ENSDARG00000004470 | prkcsh | ENSG00000130175 |
| ENSDARG00000045398 | mtrr | ENSG00000124275 |
| ENSDARG00000058158 | trim55b | ENSG00000147573 |
| ENSDARG00000100242 | dph5 | ENSG00000117543 |
| ENSDARG00000061508 | tgfbrap1 | ENSG00000135966 |
| ENSDARG00000061832 | SNTG1 | ENSG00000147481 |
| ENSDARG00000075169 | bbs1 | ENSG00000174483 |
| ENSDARG00000087780 | tiam2a | ENSG00000146426 |
| ENSDARG00000068261 | pros1 | ENSG00000184500 |
| ENSDARG00000008829 | chga | ENSG00000100604 |
| ENSDARG00000088817 | wdr59 | ENSG00000103091 |
| ENSDARG00000038300 | rnf34a | ENSG00000170633 |
| ENSDARG00000103333 | baiap2b | ENSG00000175866 |
| ENSDARG00000100515 | dusp1 | ENSG00000120129 |
| ENSDARG00000039136 | cox16 | ENSG00000133983 |
| ENSDARG00000079564 | vmhc | ENSG00000197616 |
| ENSDARG00000062758 | CR352265.1 | ENSG00000005001 |
| ENSDARG00000016481 | ptpn2a | ENSG00000175354 |
| ENSDARG00000079363 | LRIG3 | ENSG00000139263 |
| ENSDARG00000039052 | klhl40a | ENSG00000157119 |
| ENSDARG00000095947 | adkb | ENSG00000156110 |
| ENSDARG00000040627 | grik1b | ENSG00000171189 |
| ENSDARG00000077474 | PLA2R1 | ENSG00000153246 |
| ENSDARG00000100442 | cfh | ENSG00000143278 |
| ENSDARG00000077384 | dnah11 | ENSG00000105877 |
| ENSDARG00000103868 | E2F1 | ENSG00000101412 |
| ENSDARG00000019341 | gpc1a | ENSG00000063660 |
| ENSDARG00000087601 | gpr153 | ENSG00000158292 |
| ENSDARG00000055705 | f5 | ENSG00000198734 |
| ENSDARG00000068981 | glceb | ENSG00000138604 |
| ENSDARG00000098588 | gchfr | ENSG00000137880 |
| ENSDARG00000061379 | cmya5 | ENSG00000164309 |
| ENSDARG00000093235 | MPST | ENSG00000128311 |
| ENSDARG00000056768 | rprml | ENSG00000179673 |
| ENSDARG00000101057 | srsf7b | ENSG00000115875 |
| ENSDARG00000010415 | sirt4 | ENSG00000089163 |
| ENSDARG00000076730 | syt6a | ENSG00000134207 |
| ENSDARG00000042880 | si:ch211-222f23.6 | ENSG00000073008 |
| ENSDARG00000070085 | mettl6 | ENSG00000206562 |
| ENSDARG00000092507 | znf1013 | ENSG00000167394 |
| ENSDARG00000095157 | TMEM14A | ENSG00000096092 |
| ENSDARG00000054879 | six3b | ENSG00000138083 |
| ENSDARG00000103176 | nr5a1a | ENSG00000136931 |
| ENSDARG00000006604 | pvrl3b | ENSG00000177707 |
| ENSDARG00000017606 | sys1 | ENSG00000204070 |
| ENSDARG00000018327 | illr3 | ENSG00000152672 |
| ENSDARG00000016745 | slc35f6 | ENSG00000213699 |
| ENSDARG00000014274 | rfc2 | ENSG00000049541 |
| ENSDARG00000017988 | dse | ENSG00000111817 |
| ENSDARG00000040727 | tfb1m | ENSG00000029639 |
| ENSDARG00000100747 | GUCA1A (1 of 2) | ENSG00000048545 |
| ENSDARG00000061976 | sema6bb | ENSG00000167680 |
| ENSDARG00000092463 | rhbdd2 | ENSG00000005486 |
| ENSDARG00000086162 | ZNF385D | ENSG00000151789 |
| ENSDARG00000016319 | c9 | ENSG00000113600 |
| ENSDARG00000025518 | F8A2 | ENSG00000274791 |
| ENSDARG00000076176 | ptcd1 | ENSG00000106246 |
| ENSDARG00000043357 | tbl2 | ENSG00000106638 |
| ENSDARG00000013390 | ARHGAP9 | ENSG00000123329 |
| ENSDARG00000000804 | rassf6 | ENSG00000169435 |
| ENSDARG00000056615 | cybb | ENSG00000137857 |
| ENSDARG00000056099 | gtf2h5 | ENSG00000272047 |
| ENSDARG00000090963 | atp6ap1lb | ENSG00000205464 |
| ENSDARG00000054837 | zgc:136870 | ENSG00000106560 |
| ENSDARG00000079827 | srbd1 | ENSG00000068784 |
| ENSDARG00000028222 | itgb8 | ENSG00000105855 |
| ENSDARG00000009637 | zgc:73075 | ENSG00000109047 |
| ENSDARG00000032714 | gria1b | ENSG00000155511 |
| ENSDARG00000020292 | si:ch211-254n4.3 | ENSG00000184599 |
| ENSDARG00000068641 | taf10 | ENSG00000166337 |
| ENSDARG00000090401 | BCL2L14 | ENSG00000121380 |
| ENSDARG00000036063 | ppp1r11 | ENSG00000204619 |
| ENSDARG00000012763 | arl13b | ENSG00000169379 |
| ENSDARG00000076472 | ovol1 | ENSG00000083838 |
| ENSDARG00000076611 | fbxo21 | ENSG00000135108 |
| ENSDARG00000056909 | CABZ01013362.1 | ENSG00000146648 |
| ENSDARG00000102612 | slc15a4 | ENSG00000139370 |
| ENSDARG00000006074 | uck2a | ENSG00000143179 |
| ENSDARG00000070702 | s100v2 | ENSG00000189171 |
| ENSDARG00000031372 | efna2a | ENSG00000099617 |
| ENSDARG00000063212 | prmt2 | ENSG00000160310 |
| ENSDARG00000041623 | mt2 | ENSG00000087250 |
| ENSDARG00000025808 | taf5l | ENSG00000135801 |
| ENSDARG00000096989 | fam110c | ENSG00000184731 |
| ENSDARG00000015638 | gemin2 | ENSG00000092208 |
| ENSDARG00000087843 | CABZ01083501.1 | ENSG00000018236 |
| ENSDARG00000104871 | LEPROTL1 | ENSG00000104660 |
| ENSDARG00000018121 | cenpj | ENSG00000151849 |
| ENSDARG00000028586 | GPR135 | ENSG00000181619 |

| | | |
|---|---|---|
| ENSDARG00000019364 | mbip | ENSG00000151332 |
| ENSDARG00000058873 | ptpdc1b | ENSG00000158079 |
| ENSDARG00000013082 | uap1l1 | ENSG00000197355 |
| ENSDARG00000043137 | cdca8 | ENSG00000134690 |
| ENSDARG00000030156 | naglu | ENSG00000108784 |
| ENSDARG00000005924 | serpina10a | ENSG00000140093 |
| ENSDARG00000082242 | SNORA73 | ENSG00000222145 |
| ENSDARG00000087176 | sgk494a | ENSG00000167524 |
| ENSDARG00000061004 | cox11 | ENSG00000166260 |
| ENSDARG00000076755 | ap5s1 | ENSG00000125843 |
| ENSDARG00000055523 | slc22a6l | ENSG00000149452 |
| ENSDARG00000016470 | anxa5b | ENSG00000164111 |
| ENSDARG00000041538 | mrps28 | ENSG00000147586 |
| ENSDARG00000056590 | calca | ENSG00000175868 |
| ENSDARG00000060415 | arhgef28 | ENSG00000214944 |
| ENSDARG00000045025 | ift52 | ENSG00000101052 |
| ENSDARG00000077275 | crispld1a | ENSG00000121005 |
| ENSDARG00000009401 | vcanb | ENSG00000038427 |
| ENSDARG00000030239 | dennd6aa | ENSG00000174839 |
| ENSDARG00000100288 | impg2b | ENSG00000081148 |
| ENSDARG00000011926 | tpgs2 | ENSG00000134779 |
| ENSDARG00000087752 | rfwd3 | ENSG00000168411 |
| ENSDARG00000002128 | cwf19l1 | ENSG00000095485 |
| ENSDARG00000036190 | txnl4a | ENSG00000141759 |
| ENSDARG00000037706 | gss | ENSG00000100983 |
| ENSDARG00000045773 | PYURF | ENSG00000145337 |
| ENSDARG00000071084 | wipf1b | ENSG00000171475 |
| ENSDARG00000020085 | slc33a1 | ENSG00000169359 |
| ENSDARG00000089828 | C12H10orf88 | ENSG00000119965 |
| ENSDARG00000100163 | TOX4 (2 of 2) | ENSG00000092203 |
| ENSDARG00000074828 | RHOBTB2 (1 of 2 | ENSG00000008853 |
| ENSDARG00000020028 | cps1 | ENSG00000021826 |
| ENSDARG00000005989 | rgl1 | ENSG00000143344 |
| ENSDARG00000044596 | pgap2 | ENSG00000148985 |
| ENSDARG00000008859 | mylipa | ENSG00000007944 |
| ENSDARG00000073792 | man1b1a | ENSG00000177239 |
| ENSDARG00000003769 | tada2b | ENSG00000173011 |
| ENSDARG00000058964 | themis2 | ENSG00000130775 |
| ENSDARG00000058593 | sri | ENSG00000075142 |
| ENSDARG00000100260 | THEM6 (1 of 2) | ENSG00000130193 |
| ENSDARG00000005800 | ampd3a | ENSG00000133805 |
| ENSDARG00000045842 | zgc:113263 | ENSG00000092330 |
| ENSDARG00000035415 | ptger4b | ENSG00000171522 |
| ENSDARG00000075467 | shisa3 | ENSG00000178343 |
| ENSDARG00000091124 | rps19bp1 | ENSG00000187051 |
| ENSDARG00000010831 | churc1 | ENSG00000258289 |
| ENSDARG00000101814 | lpar3 | ENSG00000171517 |
| ENSDARG00000017605 | rpp40 | ENSG00000124787 |
| ENSDARG00000062594 | coq7 | ENSG00000167186 |
| ENSDARG00000102822 | pam16 | ENSG00000217930 |
| ENSDARG00000105141 | FO704635.1 | ENSG00000182261 |
| ENSDARG00000104669 | slc35c1 | ENSG00000181830 |
| ENSDARG00000071872 | zdhhc15b | ENSG00000102383 |
| ENSDARG00000062821 | slc6a15 | ENSG00000072041 |
| ENSDARG00000070434 | rhov | ENSG00000104140 |
| ENSDARG00000043566 | C9H3orf17 | ENSG00000163608 |
| ENSDARG00000090564 | PDZD4 (1 of 2) | ENSG00000067840 |
| ENSDARG00000099161 | dyx1c1 | ENSG00000256061 |
| ENSDARG00000056502 | si:ch73-334d15.4 | ENSG00000166006 |
| ENSDARG00000023583 | coq9 | ENSG00000088682 |
| ENSDARG00000075954 | serpinh1a | ENSG00000149257 |
| ENSDARG00000068104 | CABZ01041002.1 | ENSG00000204301 |
| ENSDARG00000025012 | tpi1a | ENSG00000111669 |
| ENSDARG00000078479 | mocs1 | ENSG00000124615 |
| ENSDARG00000029931 | atp13a1 | ENSG00000105726 |
| ENSDARG00000075107 | nkx2.4a | ENSG00000125816 |
| ENSDARG00000074848 | smim7 | ENSG00000214046 |
| ENSDARG00000045402 | commd3 | ENSG00000148444 |
| ENSDARG00000061195 | cnnm2a | ENSG00000148842 |
| ENSDARG00000027584 | ttpa | ENSG00000137561 |
| ENSDARG00000018065 | ntm | ENSG00000183067 |
| ENSDARG00000073999 | tapt1a | ENSG00000169762 |
| ENSDARG00000055342 | SLC16A13 | ENSG00000174327 |
| ENSDARG00000059048 | mpzl1l | ENSG00000197965 |
| ENSDARG00000017126 | ilvbl | ENSG00000105135 |
| ENSDARG00000073716 | cpxm1a | ENSG00000088882 |
| ENSDARG00000035507 | ddx31 | ENSG00000125485 |
| ENSDARG00000016835 | tcirg1a | ENSG00000110719 |
| ENSDARG00000103026 | p3h2 | ENSG00000090530 |
| ENSDARG00000069832 | sfxn4 | ENSG00000183605 |
| ENSDARG00000002037 | pfkfb2b | ENSG00000123836 |
| ENSDARG00000017703 | paqr3a | ENSG00000163291 |
| ENSDARG00000044852 | wbp2nl | ENSG00000183066 |
| ENSDARG00000014796 | wnt11r | ENSG00000085741 |
| ENSDARG00000063330 | mgat4a | ENSG00000071073 |
| ENSDARG00000100190 | MSLNL (3 of 4) | ENSG00000162006 |
| ENSDARG00000009870 | mapk8b | ENSG00000107643 |
| ENSDARG00000015662 | pla2g12b | ENSG00000138308 |
| ENSDARG00000056079 | l3mbtl2 | ENSG00000100395 |
| ENSDARG00000002847 | fndc1 | ENSG00000164694 |
| ENSDARG00000076362 | tmem260 | ENSG00000070269 |
| ENSDARG00000104242 | dcp2 | ENSG00000172795 |
| ENSDARG00000098074 | trip4 | ENSG00000103671 |
| ENSDARG00000007130 | mrto4 | ENSG00000053372 |
| ENSDARG00000012215 | umps | ENSG00000114491 |
| ENSDARG00000022845 | lias | ENSG00000121897 |
| ENSDARG00000075281 | tbc1d30 | ENSG00000111490 |
| ENSDARG00000044073 | fuca2 | ENSG00000001036 |
| ENSDARG00000052764 | chrnb3a | ENSG00000147432 |
| ENSDARG00000012745 | poc1bl | ENSG00000257594 |
| ENSDARG00000086647 | chrng | ENSG00000196811 |

| | | |
|---|---|---|
| ENSDARG00000102865 | carnmt1 | ENSG00000156017 |
| ENSDARG00000038788 | dnai1.2 | ENSG00000122735 |
| ENSDARG00000058090 | cfap57 | ENSG00000243710 |
| ENSDARG00000059175 | cep19 | ENSG00000174007 |
| ENSDARG00000086416 | med11 | ENSG00000161920 |
| ENSDARG00000055373 | sema3fb | ENSG00000001617 |
| ENSDARG00000031540 | tmem200a | ENSG00000164484 |
| ENSDARG00000035018 | thy1 | ENSG00000154096 |
| ENSDARG00000069766 | caln2 | ENSG00000183166 |
| ENSDARG00000057983 | svopl | ENSG00000157703 |
| ENSDARG00000040568 | pdzd3a | ENSG00000172367 |
| ENSDARG00000045680 | tnpo3 | ENSG00000064419 |
| ENSDARG00000104071 | C1H11orf73 | ENSG00000149196 |
| ENSDARG00000001825 | cfap43 | ENSG00000197748 |
| ENSDARG00000067912 | si:ch73-373m9.1 | ENSG00000117643 |
| ENSDARG00000059897 | ntrk2a | ENSG00000148053 |
| ENSDARG00000040295 | apoeb | ENSG00000130203 |
| ENSDARG00000016570 | prlra | ENSG00000113494 |
| ENSDARG00000055504 | CD68 | ENSG00000129226 |
| ENSDARG00000060569 | tmem229b | ENSG00000198133 |
| ENSDARG00000069996 | dnajb14 | ENSG00000164031 |
| ENSDARG00000005104 | limk2 | ENSG00000182541 |
| ENSDARG00000077532 | pign | ENSG00000197563 |
| ENSDARG00000045658 | msrb3 | ENSG00000174099 |
| ENSDARG00000075899 | adgrl2b.1 | ENSG00000117114 |
| ENSDARG00000024940 | rnf144b | ENSG00000137393 |
| ENSDARG00000030139 | sdhdb | ENSG00000204370 |
| ENSDARG00000059850 | slc35f3b | ENSG00000183780 |
| ENSDARG00000061397 | trip12 | ENSG00000153827 |
| ENSDARG00000089643 | mcama | ENSG00000076706 |
| ENSDARG00000104508 | slc10a7 | ENSG00000120519 |
| ENSDARG00000043640 | cenpn | ENSG00000166451 |
| ENSDARG00000012395 | mmp13a | ENSG00000196611 |
| ENSDARG00000071032 | pnkd | ENSG00000127838 |
| ENSDARG00000013252 | TMC5 | ENSG00000103534 |
| ENSDARG00000036764 | hax1 | ENSG00000143575 |
| ENSDARG00000057624 | cops5 | ENSG00000121022 |
| ENSDARG00000053131 | irak3 | ENSG00000090376 |
| ENSDARG00000075441 | vwa2 | ENSG00000165816 |
| ENSDARG00000086848 | atad3b | ENSG00000160072 |
| ENSDARG00000026787 | aqp7 | ENSG00000165269 |
| ENSDARG00000087206 | CCT2 | ENSG00000166226 |
| ENSDARG00000103173 | stx8 | ENSG00000170310 |
| ENSDARG00000070157 | tgm2a | ENSG00000198959 |
| ENSDARG00000043728 | srek1ip1 | ENSG00000153006 |
| ENSDARG00000101332 | uba2 | ENSG00000126261 |
| ENSDARG00000031971 | kdelc1 | ENSG00000134901 |
| ENSDARG00000075977 | prr5a | ENSG00000186654 |
| ENSDARG00000070794 | greb1 | ENSG00000196208 |
| ENSDARG00000061638 | dcakd | ENSG00000172992 |
| ENSDARG00000062549 | acmsd | ENSG00000153086 |
| ENSDARG00000079440 | coro2ba | ENSG00000103647 |
| ENSDARG00000020890 | tmod4 | ENSG00000163157 |
| ENSDARG00000010113 | ndufb8 | ENSG00000166136 |
| ENSDARG00000024314 | herpud1 | ENSG00000051108 |
| ENSDARG00000075725 | cep152 | ENSG00000103995 |
| ENSDARG00000090988 | larsa | ENSG00000133706 |
| ENSDARG00000069157 | tmem135 | ENSG00000166575 |
| ENSDARG00000042646 | ROBO3 (2 of 2) | ENSG00000154134 |
| ENSDARG00000070360 | fam212aa | ENSG00000185614 |
| ENSDARG00000056367 | mpv17l2 | ENSG00000254858 |
| ENSDARG00000006397 | laptm4a | ENSG00000068697 |
| ENSDARG00000101320 | si:dkey-50i6.5 | ENSG00000041988 |
| ENSDARG00000079632 | POLD4 | ENSG00000175482 |
| ENSDARG00000102584 | cpne7 | ENSG00000178773 |
| ENSDARG00000030775 | sybl1 | ENSG00000124333 |
| ENSDARG00000038863 | bre | ENSG00000158019 |
| ENSDARG00000055118 | mylipb | ENSG00000007944 |
| ENSDARG00000070644 | ttpal | ENSG00000124120 |
| ENSDARG00000041505 | itm2bb | ENSG00000136156 |
| ENSDARG00000063370 | sgk2a | ENSG00000101049 |
| ENSDARG00000015563 | LPIN3 | ENSG00000132793 |
| ENSDARG00000104085 | abcd3a | ENSG00000117528 |
| ENSDARG00000041576 | nudt18 | ENSG00000275074 |
| ENSDARG00000058734 | prdx1 | ENSG00000117450 |
| ENSDARG00000100352 | aglb | ENSG00000162688 |
| ENSDARG00000077537 | nudt19 | ENSG00000213965 |
| ENSDARG00000013016 | pole4 | ENSG00000115350 |
| ENSDARG00000061335 | galnt1 | ENSG00000141429 |
| ENSDARG00000071040 | smim8 | ENSG00000111850 |
| ENSDARG00000089043 | ptpn6 | ENSG00000111679 |
| ENSDARG00000052859 | chp1 | ENSG00000187446 |
| ENSDARG00000016231 | bcap29 | ENSG00000075790 |
| ENSDARG00000104254 | cpne4b | ENSG00000196353 |
| ENSDARG00000089357 | fam73b | ENSG00000148343 |
| ENSDARG00000053293 | ftr14 | ENSG00000128250 |
| ENSDARG00000004402 | elovl6 | ENSG00000170522 |
| ENSDARG00000026359 | pbld2 | ENSG00000108187 |
| ENSDARG00000043211 | ripk4 | ENSG00000183421 |
| ENSDARG00000074212 | SLC5A10 | ENSG00000154025 |
| ENSDARG00000075513 | ccdc136a | ENSG00000128596 |
| ENSDARG00000057435 | amacr | ENSG00000242110 |
| ENSDARG00000020979 | fam65c | ENSG00000042062 |
| ENSDARG00000038847 | gins3 | ENSG00000181938 |
| ENSDARG00000018562 | cope | ENSG00000105669 |
| ENSDARG00000041107 | cftr | ENSG00000001626 |
| ENSDARG00000052176 | C3H19orf66 | ENSG00000130813 |
| ENSDARG00000074866 | ptpn5 | ENSG00000110786 |
| ENSDARG00000040005 | scfd2 | ENSG00000184178 |
| ENSDARG00000097299 | FLRT2 (2 of 2) | ENSG00000185070 |
| ENSDARG00000036844 | tsen54 | ENSG00000182173 |

| | | | | |
|---|---|---|---|---|
| | | ENSDARG00000076238 | GRAMD1C | ENSG00000178075 |
| | | ENSDARG00000019877 | rbks | ENSG00000171174 |
| | | ENSDARG00000037871 | wipi2 | ENSG00000157954 |
| | | ENSDARG00000074583 | grid1a | ENSG00000182771 |
| | | ENSDARG00000100594 | sez6l | ENSG00000100095 |
| | | ENSDARG00000060153 | h6pd | ENSG00000049239 |
| | | ENSDARG00000052496 | HMHA1 (1 of 2) | ENSG00000180448 |
| | | ENSDARG00000059853 | manbal | ENSG00000101363 |
| | | ENSDARG00000099586 | CR382383.2 | ENSG00000124196 |
| | | ENSDARG00000099441 | cbx4 | ENSG00000141582 |
| | | ENSDARG00000104943 | MAD2L1BP | ENSG00000124688 |
| | | ENSDARG00000052638 | fam210b | ENSG00000124098 |
| | | ENSDARG00000098924 | suz12b | ENSG00000178691 |
