## Supplemental table 3B for "Cell type specific gene expression profiling reveals a role for the complement component C3A in neutrophil migration to tissue damage"

| Padj < 0.05 after wounding (only genes with human orthologs) | | | | | | | | |
|---|---|---|---|---|---|---|---|---|
| Neutrophils (lyz) | | | Macrophages (mpeg1) | | | Epithelium (krt4) | | |
| **Ensembl gene ID** | **Gene name** | **Human ortholog ID** | **Ensembl gene ID** | **Gene name** | **Human ortholog ID** | **Ensembl gene ID** | **Gene name** | **Human ortholog ID** |
| ENSDARG00000012694 | c3a.1 | ENSG00000125730 | | | | ENSDARG00000031119 | baiap2l1b | ENSG00000006453 |
| | | | | | | ENSDARG00000057273 | alox5a | ENSG00000012779 |
| | | | | | | ENSDARG00000010312 | cp | ENSG00000047457 |
| | | | | | | ENSDARG00000077009 | wdfy4 | ENSG00000128815 |
| | | | | | | ENSDARG00000098726 | rptor | ENSG00000141564 |
| | | | | | | ENSDARG00000063019 | panx2 | ENSG00000073150 |
| | | | | | | ENSDARG00000062101 | iffo2a | ENSG00000169991 |
| | | | | | | ENSDARG00000075673 | arhgap21b | ENSG00000107863 |
| | | | | | | ENSDARG00000100376 | amdhd2 | ENSG00000162066 |
| | | | | | | ENSDARG00000070119 | ctc1 | ENSG00000178971 |
| | | | | | | ENSDARG00000038059 | grb2b | ENSG00000177885 |
| | | | | | | ENSDARG00000074686 | mgea5 | ENSG00000198408 |
| | | | | | | ENSDARG00000038446 | nrp2b | ENSG00000118257 |
